# Exploring marine habitat restoration opportunities and blue carbon sequestration potential in the United Kingdom

**DOI:** 10.1101/2025.10.07.680237

**Authors:** Prahalad Srikanthan, Joshua Copping, Christopher Hassall

## Abstract

Nature-based solutions (NbS), including blue carbon ecosystems, are crucial to meeting Net-Zero targets. The UK’s Net-Zero 2050 strategy, alongside terrestrial measures, prioritises large-scale offshore wind development, creating opportunities to integrate marine NbS into Net-Zero plans. Consequently, aligning decarbonisation targets with conservation goals demands marine spatial planning and adaptive Marine Protected Area (MPA) networks. While estimates suggest UK marine systems store ∼244 million tonnes of organic carbon, sequestration potentials remain poorly quantified. We developed marine spatial plans for the UK and estimated carbon sequestration potentials. We divided the UK Exclusive Economic Zone into four zones (Open, Conservation, Oil, and Energy) using Marxan with Zones under two scenarios: (1) Existing MPAs locked-in, and (2) Unconstrained. Conservation Zones were defined as regions with minimal human activity. Planning incorporated 14 features: four species distribution models (SDMs), including the first UK-wide maërl SDMs, habitat suitability maps for subtidal macroalgae, and relevant stakeholder maps, such as offshore wind and oil fields. Conservation zones overlapped with >70% of current MPAs and complemented them by isolating biodiversity from anthropogenic activities, such as ports, without compromising renewable energy targets. We estimated sequestration rates of 533 - 625 kt C yr^-1^, with restoration contributing an additional 0.11 - 0.12 kt C yr^-1^ by 2040. These results provide evidence for embedding marine NbS into national strategies such as the UK Marine Strategy, while highlighting the need to address gaps in marine carbon dynamics to enable effective spatial planning and alignment with voluntary carbon markets to bridge climate finance gaps.

## Introduction

The International Union for Conservation of Nature defines Nature-based Solutions (NbS) as “*actions to protect, sustainably manage, and restore natural or modified ecosystems, that address societal challenges effectively and adaptively, simultaneously providing human well-being and biodiversity benefits*”, which can include protection, restoration and management initiatives (IUCN, 2016; O’Leary et al., 2023). As nations strive to decarbonise their economies and meet global commitments, NbS can contribute up to 25% emission reductions across transport, residential, and industrial sectors (Pan et al., 2023). Beyond carbon, NbS deliver critical ecosystem services such as flood protection, biodiversity conservation, enhanced resilience, and livelihood benefits (Chausson et al., 2020; Di Sacco et al., 2021).

The United Kingdom (UK) aims to reach Net-Zero by 2050 and clean power by 2030 by ramping up offshore wind (OW, 14.7 GW), onshore wind (15.5 GW), and solar photovoltaics (18GW) to 43-50 GW, 27-29 GW, and 45-47 GW, respectively (UK Government, 2024). Simultaneously, the UK has pledged to protect 30% of all ecosystems under the Convention on Biological Diversity (CBD), creating competing spatial demands between renewable energy expansion, biodiversity conservation, and other marine stakeholders (CBD, 2020).

The UK Exclusive Economic Zone (EEZ) supports diverse marine ecosystems, including seagrass meadows, macroalgae, maërl-forming species, kelp forests, salt marshes, seabirds, and marine mammals. The combined ecosystem services are valued at ∼£2.67 trillion, with a significant component derived from blue carbon sequestration, defined as marine and coastal carbon fluxes that can contribute to greenhouse gas (GHG) emission reductions (Küpper and Kamenos, 2018; Lovelock and Duarte, 2019; DEFRA, 2025a). UK marine systems are estimated to hold ∼244 Mt organic carbon, 98% in the top 10 cm of sediments, with ∼286,000 t C yr⁻¹ sequestered annually in coastal habitats (Burrows et al., 2024). While studies in terrestrial landscapes have shown that NbS can reduce emission by over 100% in the land sector, with a 12% reduction in food production, replicating it in marine spaces is challenging due to poor mapping, the lack of fine-scale and long-term data, and large uncertainties in ecosystem service estimates (Oestreich et al., 2020; O’Leary et al., 2023; Copping et al., 2025). For example, no UK-specific data exist for subtidal seagrass (*Zostera marina*) sequestration rates, and current values rely on global or intertidal proxies (Mcleod et al., 2011; do Amaral Camara Lima, 2020).

As marine ecosystems are increasingly threatened by climate change, overfishing, ocean acidification, and rising temperatures, it is essential to mitigate loss and conserve ecosystems (Van Der Molen et al., 2013; IPCC, 2022; Hoppit et al., 2022). While the UK EEZ contains 377 Marine Protected Areas (MPA) covering ∼38% of the seascape, several MPAs are ‘paper parks’, with over 90% of MPAs lacking site-wide regulation and frequently overlapping with high-intensity fishing, shipping, oil extraction, and wind farm development (JNCC, 2021; Greenpeace UK, 2022; Relano and Pauly, 2023). Moreover, less than 0.2% are designated as Highly Protected Marine Areas (HPMAs) with stringent regulations, raising fundamental concerns about the effectiveness of current zoning approaches in isolating conservation priorities from extractive and industrial interests (DEFRA, 2024).

Despite policy momentum, including planned trawling bans and conservation targets, key knowledge gaps persist regarding the spatial efficiency of MPAs, the feasibility of zoning strategies to reduce stakeholder conflict, and the quantifiable contribution of marine NbS to national climate targets (The Environmental Targets (Marine Protected Areas) Regulations 2023; MMO, 2025). With large overlaps between conflicting needs and little empirical evidence for blue carbon estimates in subtidal habitats, it is critical to understand whether marine spatial planning can be optimised to support conservation and energy goals. Additionally, aligning marine NbS implementation with evolving voluntary carbon markets and quantifying sequestration potentials is pivotal for supporting policy commitments and directing climate finance flows to close annual biodiversity ($342 billion) and marine ($14.6 billion) funding gaps (UNEP, 2023; Philips et al., 2025).

This study addresses these gaps by focusing on subtidal *Z. marina*, subtidal red (Rhodophyta) and brown (Phaeophyceae) macroalgae, and three maërl species (*Phymatolithon calcareum*, *Lithothamnion glaciale*, *L. corallioides*). All act as ecosystem engineers, with their three-dimensional structure providing habitat and shelter for diverse taxa (Mullan Crain and Bertness, 2006). Maërl beds support nurseries for commercially important species such as cod (*Gadus morhua*) (Kamenos et al., 2004b) and scallops (Kamenos et al., 2004a), and sustain >150 macroalgal and >500 faunal species (Birkett et al., 1998; Peña et al., 2014). They have also been observed to host twice the species richness of adjacent sites and increase community resilience (Steller et al., 2003; Bulleri et al., 2025). These habitats are highly vulnerable to bottom trawling, dredging, eutrophication, and sedimentation, leading to widespread declines across the UK (Birkett et al., 1998; Simon-Nutbrown et al., 2020). Similarly, *Z. marina* meadows provide refuges, stabilise sediments, enhance water clarity, and support food webs and fisheries (Ondiviela et al., 2014; McCloskey and Unsworth, 2015; Maxwell et al., 2017; Unsworth et al., 2019). Seagrass meadows support 39 bird species and UK’s two seahorse species, making them ecologically and culturally significant (Garrick-Maidment et al., 2010; Kollars et al., 2017). They also reduce ocean acidification and protect maërl beds from these effects (Manzello et al., 2012). Macroalgae, globally valued at ∼$1 million per km of shoreline, have several ecosystem services beyond carbon sequestration (Filbee-Dexter and Wernberg, 2018; Cotas et al., 2023; Eger et al., 2023), including positive impacts on local fisheries (Bertocci et al., 2015) and cetacean behaviour (Meynecke and Kela, 2023; Weiss et al., 2025).

In this study, we evaluated whether marine spatial planning can be optimised to spatially isolate conflicting priorities and stakeholder needs in the UK EEZ. Here, stakeholder needs refer to the spatial requirements of different marine sectors, while stakeholder uses are the specific activities they undertake. Specifically, we tested the hypothesis that spatial reconfiguration of MPAs and stakeholder use, including but not limited to oil, gas, energy, fishing, and shipping, can reduce sectoral overlap (the geographic co-occurrence of stakeholder use) while increasing the protection and restoration potential of key blue carbon habitats. To address this, we examined the technical OW potential in the UK EEZ and developed species distribution models (SDMs) for *Z. marina* and three maërl-forming species (*L. glaciale*, *L. corallioides*, *P. calcareum*). We also generated habitat suitability maps for red and brown subtidal macroalgae. This represents the first study to produce species-specific SDMs for maërl across the entire UK EEZ, addressing a critical gap and providing the first quantitative basis for incorporating maërl into marine spatial planning. Using these models and stakeholder maps, we developed two marine spatial plans to design new MPA configurations to reduce stakeholder conflict. Lastly, we estimated blue carbon sequestration potential from undisturbed habitats and habitat restoration.

## Methods

### Study area

The UK’s EEZ around the British Isles covers an area of ∼727,900 km^2^. Extending up to 200 nautical miles from the coastline, the EEZ grants the UK sovereign rights to explore, exploit, and manage natural resources, including renewable energy, fisheries, and mineral assets, as defined by the Marine and Coastal Access Act (2009) and the United Nations Convention on the Law of the Sea (UN General Assembly, 1982).

In addition to the 25+ sectors comprising the UK marine economy, estimates suggest that comprehensive mapping of the EEZ could enable activities with potential economic benefits of ∼£8.9 billion (Stebbings et al., 2020; Eunomia Research & Consulting Ltd, 2024). Key assets within the EEZ include ∼45,400 km^2^ of oil/gas fields, ∼52,490 km^2^ designated for wind farms, and commercial fisheries.

The significant potential for blue carbon sequestration, ineffective regulation in MPAs, and competing stakeholder priorities make the UK EEZ an ideal study site for evaluating MPA networks and exploring alternative marine spatial planning solutions.

### Data collection and preparation

All analyses, unless specified otherwise, were conducted using Microsoft Excel, R version 4.5.2, and the ‘sf’, ‘stars’, ‘terra’, ‘tidyverse’, ‘ggpubr’, and ‘ggplot2’ packages (Wickham, 2016; Kassambara, 2016; Pebesma, 2018; Wickham et al., 2019; Pebesma and Bivand, 2023; R Core Team, 2025; Hijmans, 2025).

We downloaded administrative boundaries, EEZ, and offshore MPA shapefiles from UKHO, (2017) and ONS UK, (2024). Spatial datasets on offshore wind farms (OWF), pipelines, oil and gas fields, shipwrecks, ports, vessel traffic density, and kinetic energy at the seabed (currents and waves) were products created by EMODnet (https://emodnet.ec.europa.eu/en/), and are owned by the EU and licensed under the CC-BY-4.0 license. We extracted 10-year averages (2010-2020) of maximum and minimum values at the seabed for salinity, temperature, nitrate, phosphate, along with surface levels of photosynthetically active radiation (PAR), and the diffuse attenuation coefficient from Bio-Oracle v3.0 using the ‘biooracler’ package (Assis et al., 2024). The diffuse attenuation coefficient is an optical property of water relating light penetration and depth (Metheniti et al., 2023). Bathymetry and wind speed at 100m were extracted from the GEBCO 2024 grid and the Global Wind Atlas, respectively (Davis et al., 2023; GEBCO Compilation Group, 2024).

Mean spring tide flow rates from the UK Renewables Atlas were used instead of Bio-Oracle v3.0 due to higher resolution (1800m vs 5600m) (ABPmer, 2008). However, since this data included flows at the water column midpoint, we estimated tidal flows 0.5m above the seabed (Equation 1). The correction was omitted for depths <0.5m. We rasterised tidal flows through universal kriging, regressing the logarithm of seabed flow velocity on depth, using the ‘gstat’ package (Gräler et al., 2016). Kriging interpolates values at unknown locations using known samples and spatial correlations (Matheron, 1963; Khan et al., 2023). The variogram was fit using the Matérn model and had a Sum of Squared Errors of 3.78e-5 (Table S1, Figure S1).

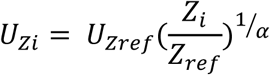

**Equation 1**. Power law relation expressing the relation between tidal current velocity and depth, where U is the tidal flow rate, i is the height above the seabed, ref is the reference height, and 𝛼 is a representation of seabed roughness, which is usually assumed to be 7 (Hill, 1991; Lee et al., 2024)

Seabed PAR was estimated using the Beer-Lambert law, incorporating depth, surface PAR, and the diffuse-attenuation coefficient (Kirk, 1994; Lund-Hansen, 2024).

### Offshore and floating technical wind potential

Technical potential represents a high-level estimate of the generation capacity that is technically achievable based solely on environmental parameters and does not account for social or economic constraints (World Bank, 2019).

We identified regions with annual average wind speeds greater than 7m/s at a 100m height in depths between 5 m and 1000 m (World Bank, 2019). Subsequently, we excluded regions that overlapped with 5 km buffer zones of ports, MPAs, pipelines in use, active oil and gas fields, shipwrecks, existing wind farms, and wind farms in the pipeline. Following this, we excluded regions with vessel densities greater than 10 hours km^-2^ month^-1^. Suitable regions less than 10 km^2^ were excluded.

We estimated potential power capacity using an average power density of 6.65 MW/km^2^ from the OWF dataset.

### Species distribution models

To obtain suitable habitats for *P. calcareum*, *L. corallioides*, *L. glaciale*, and *Z. marina,* we developed an ensemble of SDMs using maximum entropy modeling (MaxEnt) and boosted regression trees (BRT). SDMs are a widely used tool to predict species’ distributions using relationships between species presences and environmental/spatial characteristics, making them essential in informing conservation decisions (Elith and Leathwick, 2009; Elith et al., 2011; Guisan et al., 2013). We used an unweighted mean ensemble model to reduce uncertainty and bias (Araujo and New, 2007) (see supplementary methods for algorithms).

#### Presence data and environmental variables

We obtained species occurrences from the National Biodiversity Network Atlas and excluded unconfirmed/CC-BY-NC licensed observations (NBN Trust, 2025). Subsequently, using the ’ibis.iSDM’ package, we conducted spatial thinning with a 10 km radius and generated 5000 random background/pseudoabsence points (Jung, 2023).

We retained eight ecologically relevant statistically independent variables using a ±0.65 Pearson’s correlation threshold (Table 1).

**Table 1.** Species occurrence data pre and post-spatial thinning and environmental predictors used for the species distribution models

| Species | Observations | Thinned Presences | Covariates |
| --- | --- | --- | --- |
| *Zostera marina* | 5362 | 2309 | 1. Seabed Nitrate (minimum)<br>2. Seabed Salinity (minimum)<br>3. Seabed Temperature (Maximum)<br>4. Current Energy at seabed<br>5. Depth<br>6. Wave Energy at seabed<br>7. Tidal Flow at seabed<br>8. Light at seabed (minimum) |
| *Phymatolithon calcareum* | 3257 | 1754 |  |
| *Lithothamnion corallioides* | 1358 | 299 |  |
| *Lithothamnion glaciale* | 1340 | 2309 |  |

#### Model parameters and evaluation

We ran the models using the ‘maxnet’ and ‘gbm’ packages (Phillips, 2021; Ridgeway and Developers, 2024). We used linear, quadratic, and hinge feature classes for MaxEnt and set a tree complexity of 2 and learning rate of 0.01 for the BRT model. The optimum number of trees for the BRT model was estimated using the ‘gbm.step’ function from the ‘dismo’ package with a bag fraction of 0.75 (Hijmans et al., 2024). Relative feature importance was obtained from the BRT. Subsequently, we created an unweighted mean ensemble model by averaging occurrence probabilities across both models.

To assess model performance and obtain model diagnostics, we conducted a 5-fold cross-validation using the ‘mecofun’ package (Zurell, 2024). An occurrence threshold of 70% was applied to the ensemble predictions to develop a binary surface representing suitability.

### Macroalgae suitability maps

To map the potential extent of subtidal brown and red macroalgae, we developed habitat suitability layers using a threshold-based approach (see Duarte et al., (2022)) (Table S2). Additionally, we incorporated phosphate thresholds based on macroalgal tolerance to nitrogen-to-phosphorus ratios of 10:1 to 80:1 (Atkinson and Smith, 1983).

Due to the high taxonomic richness of macroalgae within the UK EEZ, we modelled macroalgae at the class level to capture broader spatial patterns. Although maërl-forming species are taxonomically Rhodophytes, we modelled their distributions separately to address the global paucity of information on maërl extents (Simon-Nutbrown et al., 2020; Tuya et al., 2023).

### Marine spatial planning

We used Marxan with Zones (MARZONE) to develop a marine spatial plan at a 25 km^2^ scale under two scenarios for the entire UK EEZ (Watts et al., 2009). MARZONE, an extension of Marxan, is a decision support tool for systematic conservation planning (Watts et al., 2009). MARZONE solves a ‘minimum set problem’ by assigning planning units (PU) to user-defined zones while meeting conservation targets at a minimum cost. Unlike Marxan, MARZONE allows the development of multiple zones with the option to assign zone-specific costs and targets, allowing practitioners to align the interests of multiple stakeholders (see Supplementary Methods).

We partitioned the EEZ into four zones: Open (OPZ), Conservation (CZ), Oil (OZ), and Energy (ENZ). The open zone comprised regions without restrictions. CZs are protected from all stakeholder activity, barring the minimal presence of OWFs due to synergies (see Discussion). OZs were included to ensure regions with oil/gas fields and pipelines were isolated from conservation regions, whereas ENZs refer to OWFs.

Scenario 1 predefined certain regions, constraining the optimisation, whereas in Scenario 2, the model was unconstrained, allowing MARZONE to design an optimal solution. Scenario 1 incorporated 14 spatial features, including six biodiversity layers comprising *L. glaciale*, *L. corallioides*, *P. calcareum*, Phaeophyceae, Rhodophyta, and *Z. marina* suitability maps, and eight stakeholder-related layers. The latter included vessel density, OW energy potential, and 5 km buffer zones around existing MPAs, pipelines, oil fields, ports, shipwrecks, and wind farms. Scenario 2 replaced existing MPAs with a spatial feature termed “protected areas,” representing each PU’s area, with a conservation target of protecting 30% of the seascape.

Feature targets differed by scenario and zone (Table 2). Biodiversity features were assigned a 30% target in alignment with the Global Biodiversity Framework (GBF) (CBD, 2020). For OW potential, a target of 0.47 was set, approximately twice the current UK OW capacity, to leverage the potential of the EEZ to accelerate renewable energy goals, facilitate international grid connectivity, and reflect the spatial availability within the EEZ to support such expansion (CCC, 2025). All other features had 100% targets. A target of 80% of MPAs in the CZ was used in Scenario 1 to ensure that 30% of the EEZ was in the CZ. Targets were considered met if 97% (Scenario 1) or 98% (Scenario 2) of each feature target was captured, as previous runs did not achieve 100% targets, allowing leeway in planning to accommodate minor spatial flexibility and data uncertainty while maintaining near-complete representation (see supplementary for model preparation and parameters).

**Table 2.** Zone targets used for Marxan with Zones in Scenarios 1 and 2

| Zone | Feature | Scenario 1 | Scenario 2 |
| --- | --- | --- | --- |
| Conservation | *Zostera marina* | 0.15 | 0.2 |
| Conservation | Rhodophyta | 0.15 | 0.2 |
| Conservation | Phaeophyceae | 0.15 | 0.2 |
| Conservation | *Phymatolithon calcareum* | 0.15 | 0.2 |
| Conservation | *Lithothamnion corallioides* | 0.15 | 0.2 |
| Conservation | *Lithothamnion glaciale* | 0.15 | 0.2 |
| Conservation | Marine Protected Areas | 0.8 | NA |
| Conservation | Protected Areas | NA | 0.3 |
| Oil | Oil Fields | 0.8 | 0.8 |
| Energy | Offshore Wind Potential | 0.47 | 0.47 |
| Energy | Wind Farms | 0.8 | 0.8 |

### Marine spatial plan analysis

To evaluate the performance and spatial characteristics across scenarios, we conducted a post-hoc analysis encompassing zone–feature contributions, selection frequency, and biodiversity representation within existing MPAs. Spatial configurations were visualised to compare final zoning solutions across scenarios. We quantified the contribution of each zone to feature-specific targets to assess how effectively zones supported biodiversity objectives and stakeholder interests.

We generated selection frequency maps for each zone based on 1000 iterations, with irreplaceable and high-priority units defined as PUs selected in 100% and ≥90% of iterations, respectively. We examined spatial similarity across scenarios using Spearman’s rank correlation (ρ) between selection frequencies in each zone.

Finally, to examine the prevalence of spatial inefficiencies or regions within the CZ that remain under the influence of conflicting activities, such as proximity to ports, we quantified the proportion of each biodiversity feature within the CZ that could be designated as undisturbed habitat (Wilson et al., 2025). Undisturbed habitat was defined as regions free from human activities using buffers of 20 km for major ports and 10 km for oil/gas fields, OWFs, and regions with high vessel density (Martino, 2001).

### Blue carbon sequestration potentials

Using Monte Carlo simulations, we estimated carbon sequestration potentials for each biodiversity feature across scenarios until 2040 (n = 1,000). To avoid double-counting, Rhodophyta estimates excluded overlaps with Phaeophyceae distributions. This represented an upper bound of potential sequestration by assuming that all undisturbed habitats constituted ecologically viable blue carbon storage habitats.

We used a global average of 330 g C m^-2^ yr^-1^ for maërl organic carbon sequestration due to limited species-specific data (Van Der Heijden and Kamenos, 2015; Cunningham and Hunt, 2023). Inorganic carbon sequestration from calcification was excluded due to CO_2_ release during the process (Frankignoulle et al., 1994; Turrell et al., 2023). Due to the lack of regional data and high variability in global estimates, we used UK-based estimates from intertidal seagrass sequestration rates for *Z. marina* (Mcleod et al., 2011; do Amaral Camara Lima, 2020) (Table 3). We simulated sequestration values for *Z. marina* and maërl using normal distributions, with a 10% standard deviation applied to maërl.

**Table 3.** Monte Carlo simulation parameters. Sequestration for Rhodophyta and Phaeophyceae represents Net Primary Productivity (see Methods). Mean maërl project sizes were estimated as half the global median (Danovaro et al., 2025). SD - Standard Deviation. Mean restoration size is on the lognormal scale for *Zostera marina*, Rhodophyta, and Phaeophyceae

| Species | Sequestration<br>(g C m<sup>-2</sup> yr<sup>-1</sup>) | SD | Mean<br>restoration<br>size (km<sup>2</sup>) | SD | #<br>projects/year | Success<br>% | Logistic<br>growth<br>rate | Logistic<br>midpoint |
| --- | --- | --- | --- | --- | --- | --- | --- | --- |
| *Zostera marina* | 67.91 | 32.39 | -8.72541 | 3.30 | 5 | 56 | 1.17 | 2.5 |
| Rhodophyta | 105 | 130 | -9.15014 | 4.04 | 5 | 60 | 1.5 | 2 |
| Phaeophyceae | 362.2859 | 632 | -9.15014 | 4.04 | 5 | 60 | 1.5 | 2 |
| *Phymatolithon calcareum* | 330 | 33 | 0.02 | 0.006 | 5 | 64 | 0.058 | 51 |
| *Lithothamnion corallioides* | 330 | 33 | 0.02 | 0.006 | 5 | 64 | 0.058 | 51 |
| *Lithothamnion glaciale* | 330 | 33 | 0.02 | 0.006 | 5 | 64 | 0.058 | 51 |

Phaeophyceae sequestration estimates were derived from net primary productivity (NPP) data across Spain, Iceland, the UK, Germany, Norway, Greenland, and Denmark, with outliers (bottom and top 5%) removed (Pessarrodona et al., 2021) (Table 3). Rhodophyta NPP values were sourced from Duarte et al., (2022) (Table 3). We used gamma distributions, truncated at observed minima and maxima, to simulate NPP for macroalgal groups (Pessarrodona et al., 2021). Macroalgal sequestration was assumed to represent 1-20% of NPP, sampled from a uniform distribution (Krause-Jensen and Duarte, 2016; Eger et al., 2023).

Lastly, we simulated restoration projects until 2040. Using data from the Global Marine Restoration Database, we modelled five projects per feature per year to reflect an ideal scenario, which exceeds recent UK restoration rates of 27 projects from 2015–2022 (UNEP-WCMC, 2022). We modelled annual project sizes for seagrass and macroalgae using a lognormal distribution from available data (Eger et al., 2023; Danovaro et al., 2025) (Table 3). Mean maërl project size was half the global median (0.04 ha) due to large knowledge gaps in maërl restoration, which are likely to result in less ambitious efforts (Tuya et al., 2023; Danovaro et al., 2025). Project growth was simulated using a logistic function, assuming 95% of sequestration potential is achieved at maturity. Logistic midpoints were assigned based on species-specific maturation rates and project delays. Restoration outcomes were multiplied by project success rates to obtain expected sequestration estimates (Danovaro et al., 2025). Success rates, logistic midpoints, and maërl project size were normally distributed with standard deviations equal to 5% of the mean.

We computed summary statistics using the ‘DescTools’ package (Signorell, 2014). Ninety-five percent confidence intervals (CI) were estimated for means and medians, with bootstrapped intervals applied to macroalgae and total sequestration due to non-normality. To assess differences between scenarios in 2026 and 2040, we applied Mann-Whitney U tests with a Benjamini-Hochberg correction for multiple comparisons.

Species-level restoration potential was quantified as the difference in median sequestration between 2026 and 2040 across simulations, with uncertainty estimated using bootstrapped 95% CIs.

## Results

### Offshore and floating wind energy potential

Thirty percent (219,524 km^2^) of the EEZ was identified as technically suitable for offshore and floating wind energy development, corresponding to an estimated 1,459.83 GW of technical potential (Figure 2).

**Figure 1.**
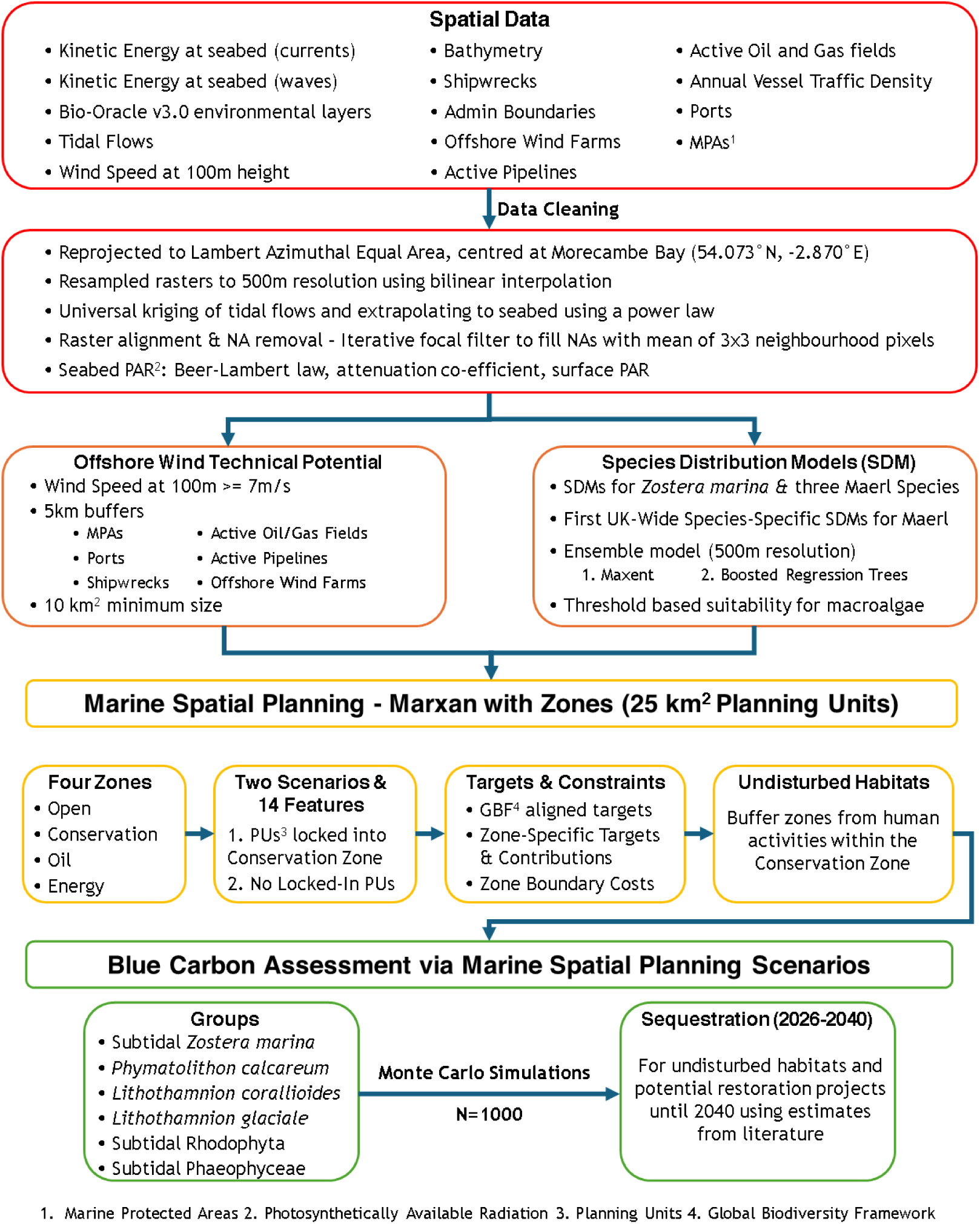
Methodology workflow for marine spatial planning and blue carbon assessment in UK waters. Outline of steps taken, from spatial data to marine spatial planning, followed by estimation of blue carbon sequestration potentials

**Figure 2.**
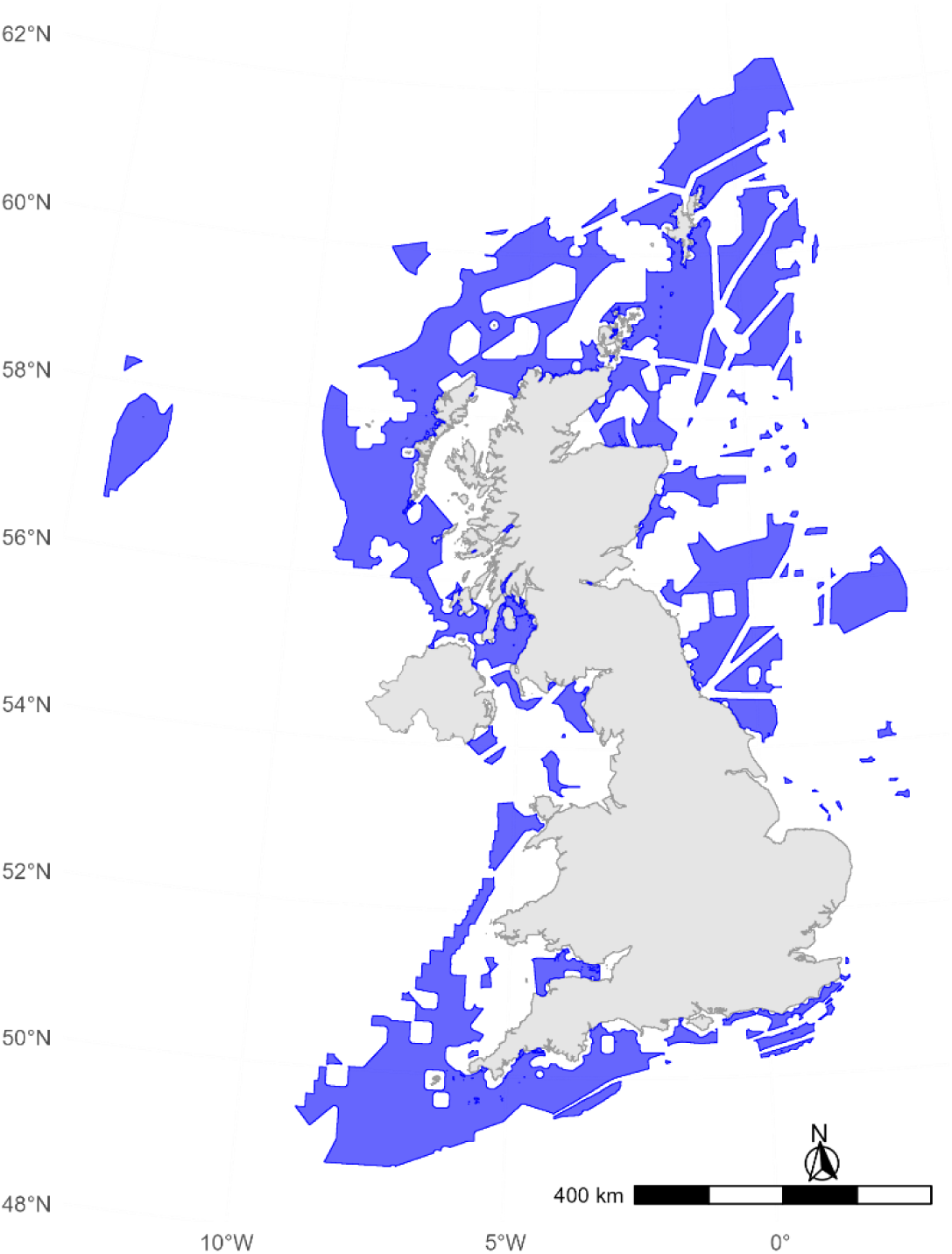
Map of offshore wind energy technical potential across the UK Exclusive Economic Zone at a 500m resolution. Areas in blue indicate locations with suitable technical potential for offshore and floating wind development. Technical potential represents a high-level estimate of the generation capacity that is technically achievable based solely on environmental parameters (World Bank, 2019)

**Figure 3.**
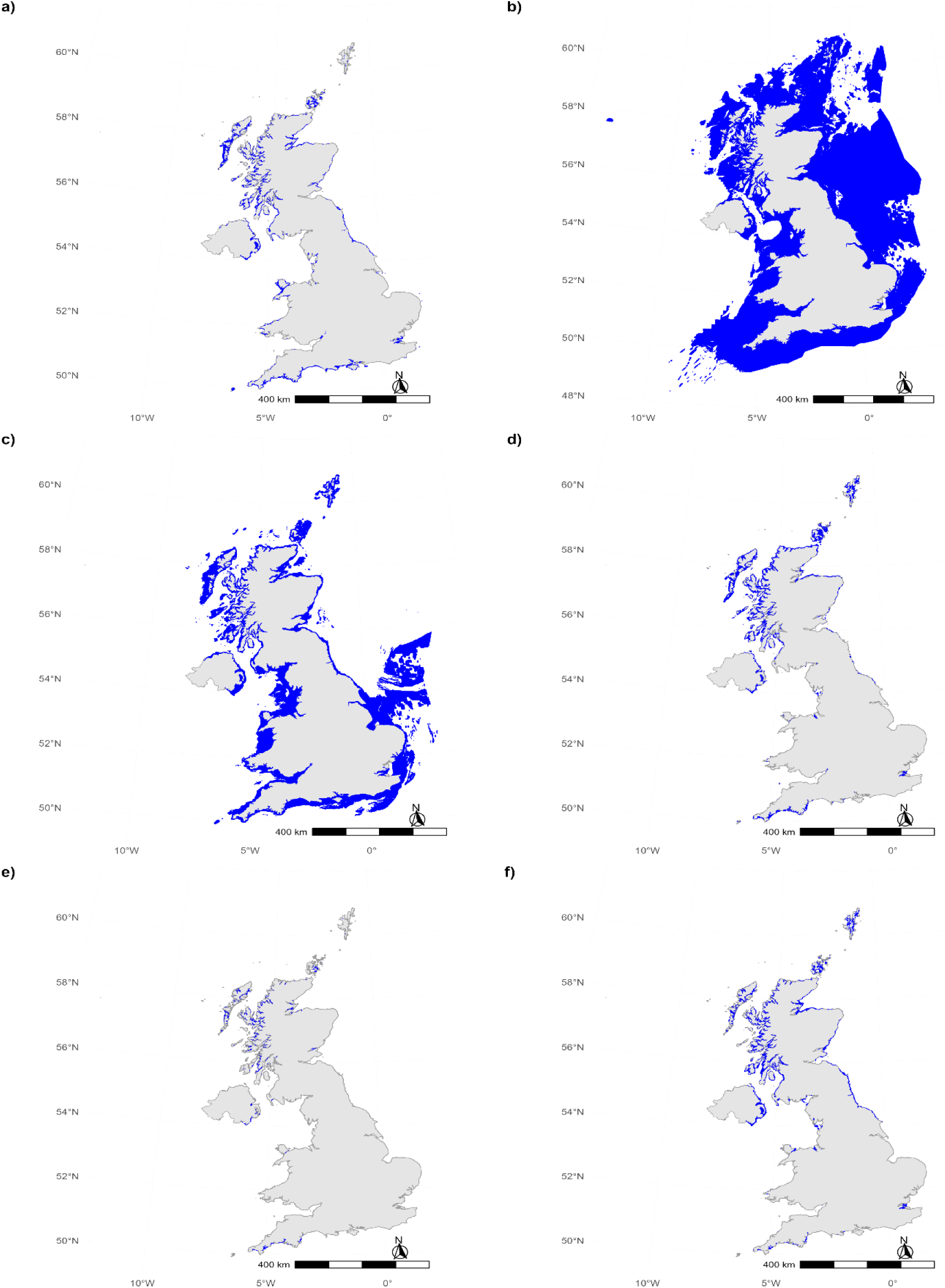
Species distribution models and habitat suitability maps in the UK Exclusive Economic Zone for (a) *Zostera marina*, (b) Rhodophyta (red macroalgae), (c) Phaeophyceae (brown macroalgae), (d) *Phymatolithon calcareum*, (e) *Lithothamnion corallioides*, and (f) *Lithothamnion glaciale*. Blue indicates suitable habitat

### Habitat Suitability

SDMs exhibited high predictive performance across algorithms, with AUCs>0.97 and TSS>0.85 (Table 4).

**Table 4.** Model evaluation metrics for four species distribution models obtained from 5-fold cross-validation of Maxent, Boosted Regression Trees (BRT), and an Ensemble model. AUC - Area Under the Curve, TSS - True Skill Statistic, PCC - Percent Correctly Classified, and deviance explained (D²)

| Species | Model | AUC | TSS | Kappa | Sensitivity | Specificity | PCC | D<sup>2</sup> |
| --- | --- | --- | --- | --- | --- | --- | --- | --- |
| *Zostera marina* | Maxent | 0.985 | 0.894 | 0.808 | 0.922 | 0.972 | 0.968 | 0.635 |
|  | BRT | 0.992 | 0.946 | 0.832 | 0.976 | 0.971 | 0.971 | 0.811 |
|  | Ensemble | 0.993 | 0.940 | 0.838 | 0.967 | 0.973 | 0.972 | 0.756 |
| *Phymatolithon calcaereum* | Maxent | 0.977 | 0.854 | 0.584 | 0.959 | 0.895 | 0.901 | 0.598 |
|  | BRT | 0.988 | 0.910 | 0.781 | 0.951 | 0.959 | 0.958 | 0.727 |
|  | Ensemble | 0.986 | 0.902 | 0.757 | 0.949 | 0.953 | 0.953 | 0.697 |
| *Lithothamnion corallioides* | Maxent | 0.987 | 0.922 | 0.321 | 0.964 | 0.958 | 0.958 | 0.659 |
|  | BRT | 0.981 | 0.899 | 0.649 | 0.909 | 0.990 | 0.989 | 0.673 |
|  | Ensemble | 0.989 | 0.932 | 0.388 | 0.964 | 0.968 | 0.968 | 0.715 |
| *Lithothamnion glaciale* | Maxent | 0.984 | 0.881 | 0.651 | 0.949 | 0.932 | 0.934 | 0.640 |
|  | BRT | 0.990 | 0.925 | 0.752 | 0.968 | 0.956 | 0.957 | 0.742 |
|  | Ensemble | 0.990 | 0.918 | 0.752 | 0.961 | 0.957 | 0.957 | 0.729 |

**Table 5.** Marxan with Zones output of the best solutions for Scenarios 1 and 2. MPM refers to the minimum proportion met for the feature. Overall targets were considered to be met if >97% or 98% were met for Scenarios 1 and 2, respectively. Zone-specific targets were met for all features

| Feature | Scenario 1 |  | Scenario 2 |  |
| --- | --- | --- | --- | --- |
|  | Target Met | MPM | Target Met | MPM |
| Protected Areas | NA | NA | Yes | 1 |
| Marine Protected Areas | No | 0.932 | NA | NA |
| Shipwrecks | Yes | 1 | Yes | 0.996 |
| Vessels | Yes | 1 | Yes | 1 |
| Ports | No | 0.741 | Yes | 0.998 |
| Pipelines | Yes | 1 | Yes | 1 |
| Oil Fields | Yes | 0.982 | Yes | 0.996 |
| Wind Farms | Yes | 0.974 | Yes | 0.986 |
| Offshore Wind Potential | Yes | 1 | Yes | 1 |
| *Lithothamnion glaciale* | Yes | 1 | Yes | 1 |
| *Lithothamnion corallioides* | Yes | 1 | Yes | 1 |
| *Phymatolithon calcareum* | Yes | 1 | Yes | 1 |
| Phaeophyceae | Yes | 1 | Yes | 1 |
| Rhodophyta | Yes | 1 | Yes | 1 |
| *Zostera marina* | Yes | 1 | Yes | 1 |

Light at seabed was the most important predictor for all species (Figure S2). For *P. calcareum* and *L. glaciale*, seabed temperature and tidal flow were the next most influential variables, whereas wave energy and current velocity contributed most to *L. corallioides* model performance. Depth and seabed temperature were the next strongest predictors for *Z. marina*.

Using a 70% occurrence threshold on ensemble predictions, *P. calcareum* had the largest habitat extent (4,116.48 km^2^), followed by *L. glaciale* (2,883.87 km^2^), *Z. marina* (2,788.41 km^2^), and *L. corallioides* (400.43 km^2^). Predicted habitat suitability based on environmental thresholds for subtidal brown and red macroalgae was 80,695.78 km^2^ and 342,192.83 km^2^, respectively.

### Marine Spatial Planning

#### MARZONE

The CZ in Scenario 1 covered 40% of the EEZ and 84% of the current MPA network compared to 37% and 74% in Scenario 2. In Scenario 1, scores ranged from 3454 to 71,899 (median: 70,430). Scenario 2 resulted in substantially higher scores, ranging from 641,323 to 705,593 (median: 699,275). PU allocation differed across zones, with 12,604 PUs allocated to the CZ in Scenario 1 compared to 11,528 in Scenario 2. Scenario 2 allocated more PUs to the ENZ (7554 vs. 7774) and OZ (3610 vs. 3689).

Scenario 1 failed to targets for MPAs and ports, with only 93.2% of MPAs captured and 74.1% of port areas retained. These shortfalls resulted from 20,515.65 km^2^ of MPAs falling within the OZ and 433 km^2^ of ports within CZs (Figure 5). In contrast, Scenario 2 met all targets. CZs in Scenario 1 had larger overlaps with ports, oil fields, wind farms, and regions with high vessel density than Scenario 2 (Table S5a,b).

**Figure 4.**
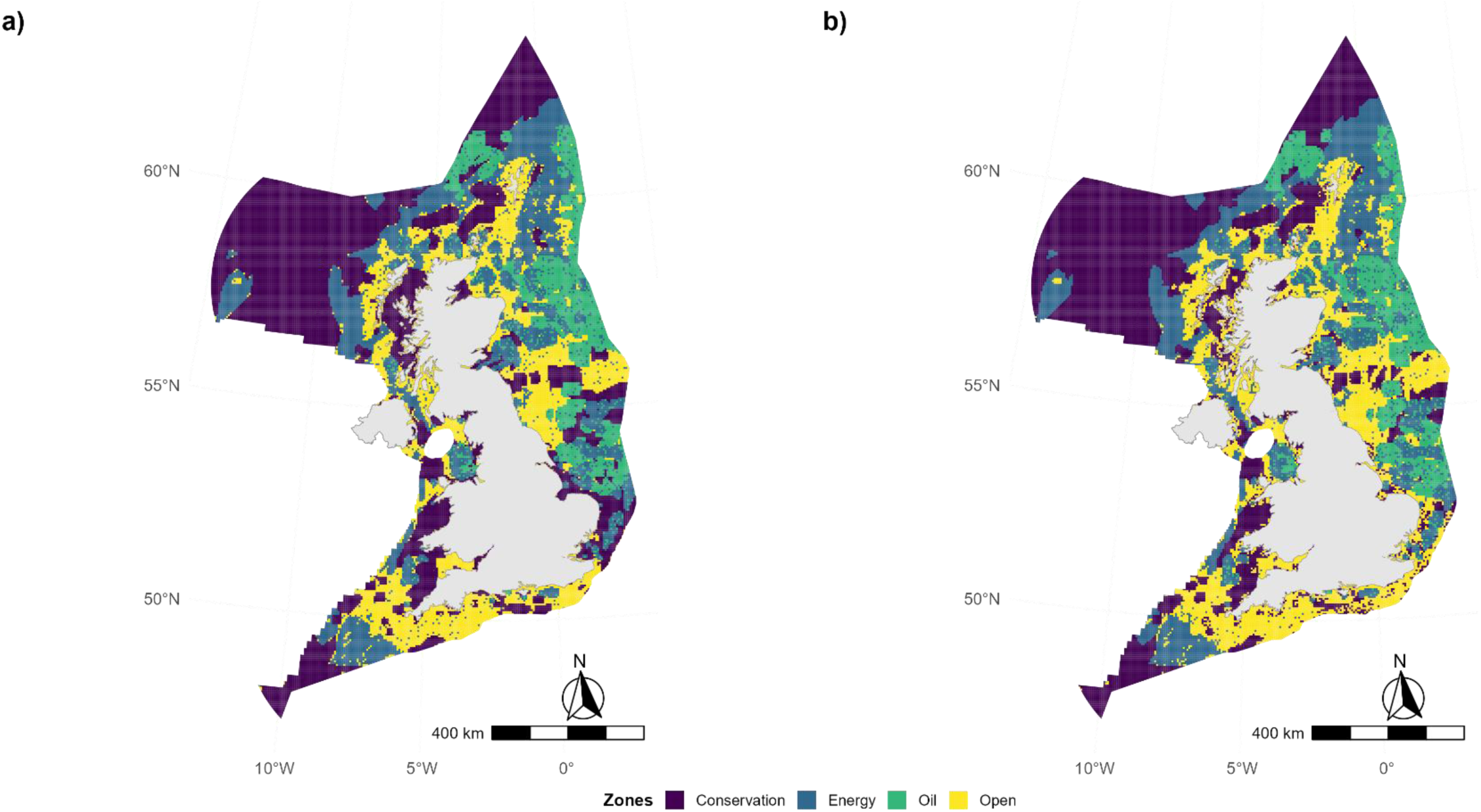
Marxan with Zones marine spatial plans for a) Scenario 1 - with Marine Protected Areas (MPA) locked in and 15% biodiversity conservation target in conservation zones (CZ) b) Scenario 2 - with no locked in units and 20% biodiversity conservation target in CZs. Both scenarios were also required to accommodate 30% of the Exclusive Economic Zone in CZs (CBD, 2020).

**Figure 5.**
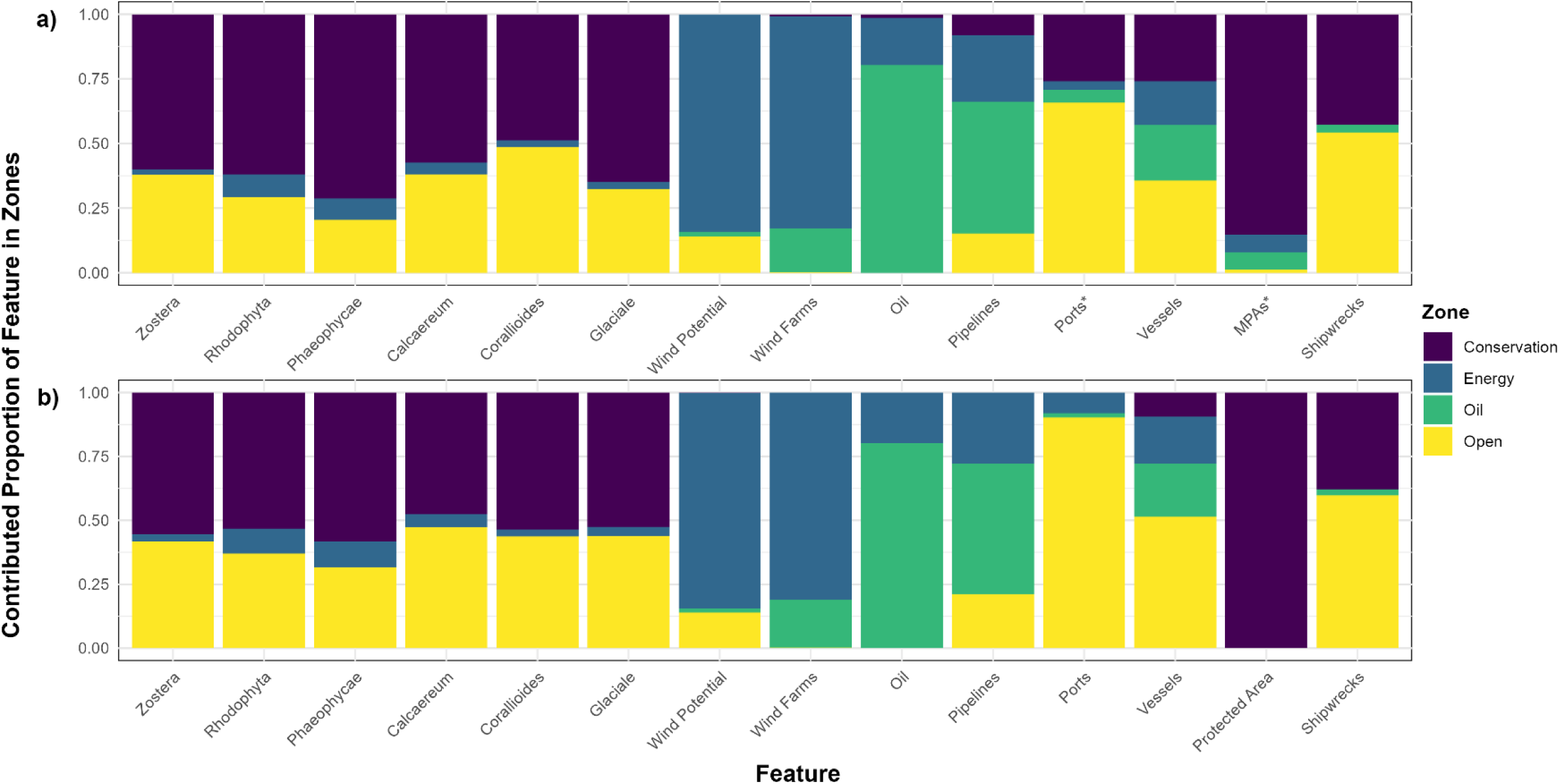
Proportion of contributed amounts per zone to each feature under (a) Scenario 1 and (b) Scenario 2. Stacked bars show the relative contribution of each management zone to overall feature targets. Asterisks (*) indicate features for which targets were not met. For Marine Protected Areas (MPA), Oil, and Ports, bars represent the full zonal breakdown and not contributed amounts

Each biodiversity feature had a total conservation target of 30% of its extent, with the CZ target set at 15% in Scenario 1 and 20% in Scenario 2 (Table 2). All biodiversity features exceeded CZ targets across scenarios, except Rhodophyta in Scenario 2 (Table S4a,b, Figure 6a). Phaeophyceae exceeded its CZ target by 15,933.5 km^2^ and 2921 km^2^ in Scenarios 1 and 2. *L. glaciale* surpassed its target by 557.21 km^2^ and 117.9 km^2^, while *L. corallioides* exceeded its target by 26.6 km^2^ and 20.1 km^2^ in Scenarios 1 and 2, respectively. *Z. marina* was allocated 400.5 km^2^ and 168 km^2^ above its target in Scenarios 1 and 2, respectively. Rhodophyta exceeded its target by 37,677.42 km^2^ in Scenario 1 but met it exactly in Scenario 2.

**Figure 6.**
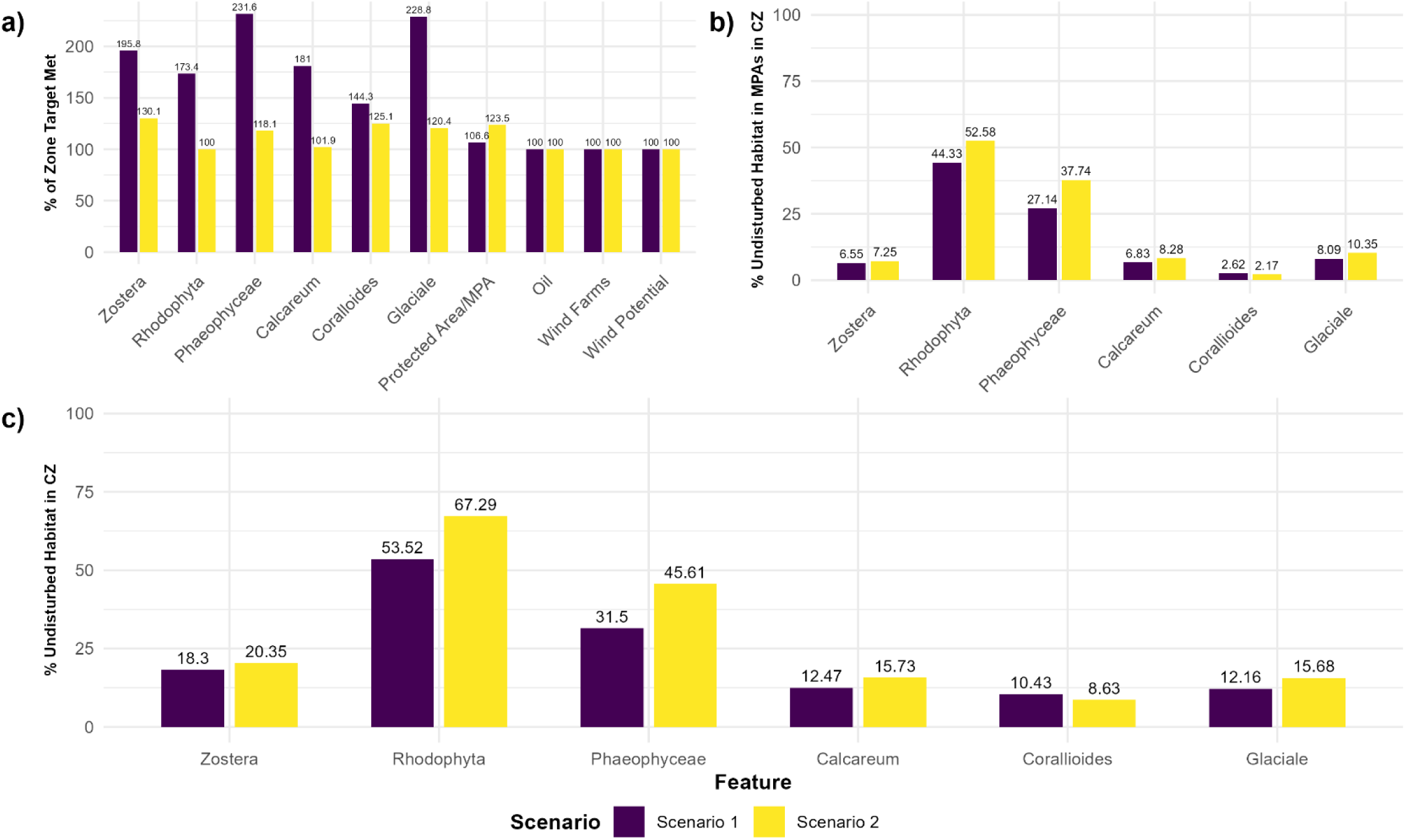
(a) Percentage of zone targets met for each feature (see Table 3) (b) Proportion of suitable habitat that is undisturbed within the Conservation Zone that overlaps with MPAs (c) Proportion of suitable habitat that is undisturbed habitat within the Conservation Zone. CZ – Conservation Zone; MPA – Marine Protected Area, Zostera - *Zostera marina*, Calcareum - *Phymatolithon calcareum*, Corallioides - *Lithothamnion corallioides*, Glaciale - *Lithothamnion glaciale*.

#### Selection Frequencies

Scenario 1 consistently yielded higher proportions of high-priority PUs (selected in ≥90% of iterations, indicating high importance) and irreplaceable PUs (selected in 100% of iterations, indicating essential inclusion) (Table S7, Figure S3). In the CZ of Scenario 1, 99.8% of PUs were irreplaceable or high-priority compared to 76.9% in Scenario 2. The OZ, while having the lowest proportion of irreplaceable PUs (<1% across scenarios), showed greater high-priority representation under Scenario 1 (52.3%) than under Scenario 2 (37.34%). Similarly, the ENZ maintained ∼27% high-priority and irreplaceable PUs across scenarios.

Selection frequencies were strongly correlated across scenarios within each zone, with Spearman’s rank correlation coefficients exceeding 0.87 for all zones (ρ = 0.878-0.963, *p* < 0.001) (Table S8).

#### Undisturbed habitats within the Conservation Zone

The percentage of undisturbed habitat in the CZ varied across species (Figure 6b,c). While Scenario 2 resulted in larger percentages of suitable area in undisturbed habitat, overall areas remained similar across scenarios (Table S9). Despite substantial overlap between the CZ and existing MPAs, undisturbed areas for coastal species occurred predominantly outside the current MPA network across scenarios (Figure 6b,c). Phaeophyceae and Rhodophyta exhibited the highest proportions of undisturbed habitat, whereas *L. corallioides* had the lowest extent of undisturbed suitable area within the CZ.

### Blue Carbon Sequestration

Total carbon sequestration was consistently higher under Scenario 1 (Figure 7, Table S10). In 2026, Scenario 2 yielded a median total of 573,049.70 t C yr⁻¹ [95% CI: 533,206.6 - 610,521.2], compared to 587,123 t C yr⁻¹ [545,460.4 - 625,558.4] under Scenario 1. By 2040, medians showed a moderate increase, reaching 574,790.90 t C yr⁻¹ [533,290.5 - 6,11,044.5] and 589,002.60 t C yr⁻¹ [545,568.7 - 625,732.1] under Scenarios 2 and 1, respectively. Mean values reached 778,557.30 t C yr⁻¹ [740,114.79 - 815,633.9] in 2026 and 778,804.30 t C yr⁻¹ [740,891.45 - 817,866.23] in 2040, compared to 758,614.30 t C yr⁻¹ [722,564.4 - 795,742.37] and 758,861.30 t C yr⁻¹ [719,510.29 - 796,259.36] under Scenario 1 and 2, respectively.

**Figure 7.**
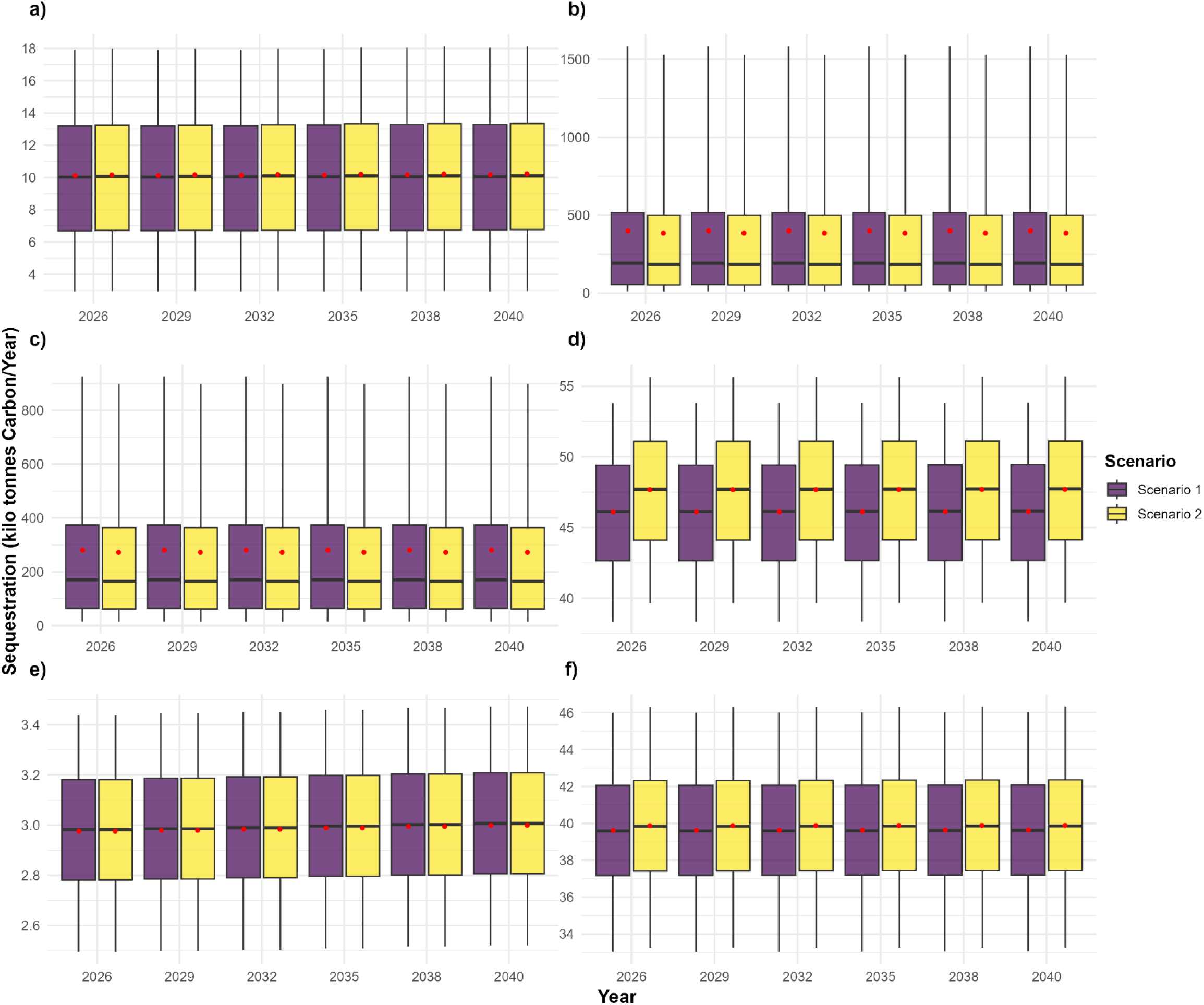
Projected annual blue carbon sequestration from 2026 to 2040 for (a) *Zostera marina*, (b) Rhodophyta (red macroalgae), (c) Phaeophyceae (brown macroalgae), (d) *Phymatolithon calcareum*, (e) *Lithothamnion corallioides*, and (f) *Lithothamnion glaciale* under Scenario 1 (purple) and Scenario 2 (yellow). Horizontal lines represent the median and whiskers extend to the 5th and 95th percentiles. Red dots indicate the mean of all simulation runs.

At the species level, red and brown macroalgae exhibited the largest sequestration potentials across scenarios and years, with median values under Scenario 1 reaching 190,725.80 t C yr⁻¹ [168,620.32 - 209,657.33] and 170,400.20 t C yr⁻¹ [152,726.27 - 186,559.03], respectively, in 2026. These values remained consistent in 2040, with Rhodophyta sequestering 1,90,728.10 t C yr⁻¹ [1,74,576.88 - 2,17,063.08] and Phaeophyceae sequestering 170,403.50 t C yr⁻¹ [152,805.41 - 186,563.12]. *Z. marina* showed moderate sequestration potential, with medians of 10,026.80 t C yr⁻¹ [9557.24 - 10,277.52] in 2026 and 10,051.70 t C yr⁻¹ [9677.9 - 10,300.54] in 2040 under Scenario 1. *L. corallioides* had the lowest sequestration estimates under both scenarios, with a median of 2982.10 t C yr⁻¹ [2957.23 - 2999.88] in 2026 and 3006.70 t C yr⁻¹ [2979.3 - 3026.24] in 2040 under Scenario 1.

In both 2026 and 2040, we did not find significant differences in sequestration rates across scenarios, except for *P. calcareum,* where Scenario 2 resulted in higher sequestration rates (Mann-Whitney U: U=5.90e+05, p_adj <0.01, Table S11).

Restoration projects, across scenarios, were estimated to sequester 115.0 t C yr⁻¹ [110.9-122.3] in 2040 (Table S12). Maërl restoration contributed the largest gains, with additional sequestrations of 23 t C yr⁻¹ [22-24] (Figure 8, Table S11). *Z. marina* restoration resulted in additional sequestrations of 9.6 t C yr⁻¹ [8.2-10.8], while Phaeophyceae showed smaller gains (7.6 t C yr⁻¹ [6.2-9.2]). Rhodophyta contributed the lowest sequestration gains at 1.45 t C yr⁻¹ [1.19-1.76].

**Figure 8.**
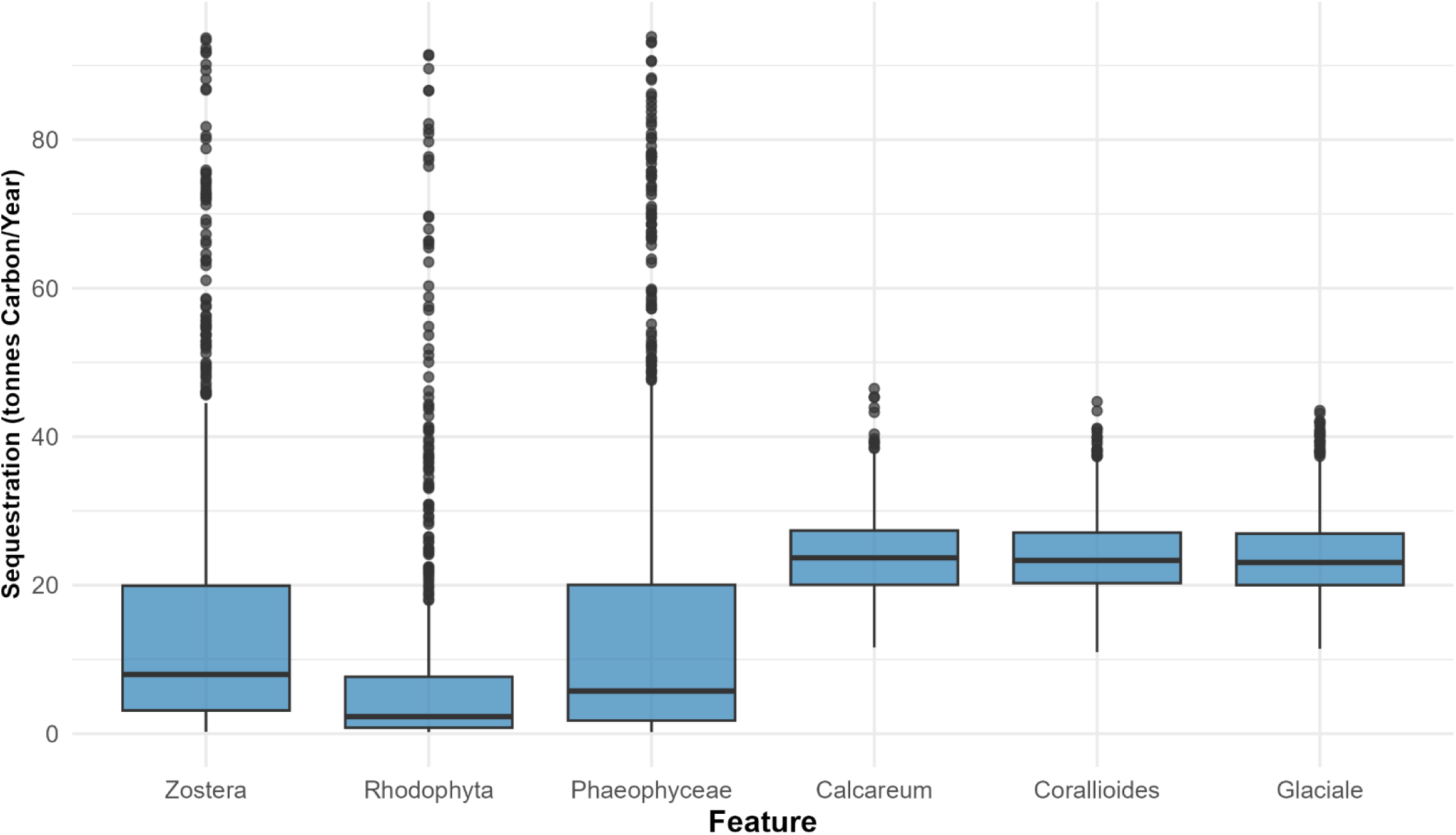
Carbon sequestered (t C yr^-1^) in 2040 by restoration projects developed from 2026-2040. Zostera - *Zostera marina*, Calcareum - *Phymatolithon calcareum*, Corallioides - *Lithothamnion corallioides*, Glaciale *- Lithothamnion glaciale*. Horizontal lines represent the median and whiskers extend to the 5th and 95th percentiles

## Discussion

International agreements mandating conservation targets complement net-zero ambitions by leveraging the substantial carbon sequestration potential offered by NbS. However, achieving these targets requires reconciling the competing demands of conservation, renewable energy, and diverse stakeholder priorities through top-down planning and effective implementation. In marine contexts, this often involves spatial zoning informed by stakeholder-driven marine spatial planning to balance conservation with human activities.

We propose a marginally new MPA network to drastically reduce overlap of biodiversity with conflicting interests, such as ports, without compromising stakeholder interests and national commitments. Importantly, our analysis quantified significant restoration potentials, revealing blue carbon sequestration capacities estimated between 533,206.6 - 625,558.4 tonnes of carbon annually, amounting to 0.6% of the UK’s annual emissions, along with a significant potential for additional OW projects, highlighting the important role the UK can play in meeting global net-zero targets.

### SDMs and habitat suitability

#### SDMS

We developed four ensemble SDMs with a 70% occurrence probability threshold representing the first UK-wide quantification of suitable maërl habitats. Our models outperformed earlier approaches, incorporating larger presence datasets and producing more conservative estimates across species (Simon-Nutbrown et al., 2020; Castle et al., 2022).

Compared to previous estimates of suitable *Z. marina* habitat of 4,194.1 km^2^ (300m resolution), we identified 2,788.41 km^2^ (500m resolution) within the UK EEZ, which exceeds mapped distributions highlighting historical declines (Green et al., 2021; Castle et al., 2022). Our model had a higher AUC (0.993 vs 0.941) and used more presence points (2309 vs 256). Consistent with previous studies, minimum light at seabed (35%) and depth (28%) were the most important parameters (Bekkby et al., 2008; Castle et al., 2022). Bottom temperature (14%), the third most important variable, has been linked to increasing seagrass mortalities (Wilson and Lotze, 2019). Interestingly, despite seagrasses facing threats from increased nitrogen concentrations, we found minimum seabed nitrate was the 4^th^ most important variable (5%) (Jones et al., 2018). Given the strong correlation between minimum and maximum nitrate (r = 0.94), the nutrient-enriched status of UK waters, and the role of light limitation at the seabed, we infer that habitat suitability reflects an interaction between indirect nutrient-driven light attenuation due to algal blooms and direct impacts of enrichment due to carbon imbalances arising from increased nitrogen assimilation (Duarte, 1995; Burkholder et al., 2007; Jones et al., 2018). Including nitrate as a predictor and differences in resolution likely contribute to differences in the estimates relative to Castle et al., (2022).

Our maërl SDMs identified ∼7,400 km^2^ of suitable habitat in the UK, spread across *P. calcareum* (4,116.48 km^2^), followed by *L. glaciale* (2,883.87 km^2^), and *L. corallioides* (400.43 km^2^). A previous study identified 7130 km^2^ of suitable habitat in Scotland, with an AUC of 0.959 compared to our average AUC of 0.988 (Simon-Nutbrown et al., 2020). Unlike our models, Simon-Nutbrown et al., (2020) did not separate species and used only 1,430 occurrence records, fewer than our species-specific datasets, barring *L. corallioides*, supporting the higher accuracy of our estimates.

For both *P. calcareum* and *L. glaciale*, minimum light at the seabed, maximum seabed temperature, and tidal flow were the top predictors. In contrast, previous models highlighted depth, likely reflecting differences in species separation and presence data (Simon-Nutbrown et al., 2020). Tidal regimes influence maërl distributions by impacting irradiance and mobility (Birkett et al., 1998). For *L. corallioides*, wave energy and current velocity at the seabed were the second and third most important predictors, aligning with their roles in influencing sediment burial and smothering risk (Perry and Jackson, 2017). Wave exposure has also been observed to impact species composition, with *L. corallioides* preferring sheltered regions with lower exposure (Bosence, 1976; Birkett et al., 1998). Differences in the importance of flow-and energy-related predictors may reflect smaller sample sizes for *L. corallioides* and species-specific morphological traits (Melbourne et al., 2018). Projected climate scenarios show significant habitat contraction; however, species-specific thermal tolerances suggest differential responses to warming, highlighting an urgent need for large-scale conservation measures (Kamenos and Law, 2010; Pardo et al., 2014; Simon-Nutbrown et al., 2020).

While our models showed high predictive ability (AUCs > 0.977, TSS > 0.854), they must be cautiously interpreted due to potential misrepresentations of species prevalences resulting from the lack of true absence data (Allouche et al., 2006; Jiménez-Valverde, 2012; Leroy et al., 2018). Without true absence data, we recommend that future studies incorporate similarity metrics or include species prevalence as an additional parameter (Leroy et al., 2018). Despite these limitations, our SDMs provide the most conservative and spatially comprehensive estimates of suitable maërl and seagrass habitats, offering a foundation for prioritising conservation and restoration efforts.

#### Macroalgae habitat suitability

Using environmental thresholds, we estimated the upper limits of suitable habitat for subtidal brown and red macroalgae within the UK EEZ as 80,695.78 km^2^ and 342,192.83 km^2^, respectively. Global estimates for subtidal brown and red algae range from 1.43-2.50 million km² and 2.98-6.69 million km^2^, respectively, with UK-specific estimates placing suitable subtidal brown algal habitat between 8000 km^2^ - 30,000 km^2^ (Yesson et al., 2015; Smale et al., 2016; Duarte et al., 2022; Burrows et al., 2024). With no UK-specific estimates for subtidal Rhodophyta, our estimates address this gap and refine global approaches by incorporating phosphate thresholds alongside those proposed by Duarte et al., (2022).

### Marine Spatial Planning

We compared zoning plans across two scenarios: Scenario 1 locked in existing MPAs, while Scenario 2 allowed MARZONE to determine optimal zoning without predefined constraints. Scenario 1 classified 40% of the EEZ as CZ, compared to 37% in Scenario 2, both surpassing GBF targets (CBD, 2020). The high overlap of the CZ (84%, 74%) with the current MPA network and high correlation between scenarios indicate that the MPA network is effective at capturing biodiversity.

On one hand, the UK has the largest MPA network in the world, covering ∼38% of the EEZ, and has committed to ambitious restoration targets, including the Environmental Targets (MPAs) Regulations 2023, which aim to ensure at least 70% of protected features are in favourable condition by 2043, and the OSPAR Convention, which promotes connectivity between North-East Atlantic MPAs (JNCC, 2021; DEFRA, 2024). On the other hand, over 90% of these MPAs lack site-wide regulation, with some being considered as ‘paper parks’ or MPAs in name only (Greenpeace UK, 2022; Relano and Pauly, 2023). This dichotomy is reflected in our results, which show significant overlaps with oil, wind farms, ports, and the CZ in Scenario 1. The low proportion of undisturbed coastal habitats within MPAs further highlights the limitations of the current network.

Despite meeting slightly lower conservation targets than Scenario 1, Scenario 2 addresses these critical issues by efficiently separating conflicting spatial priorities across stakeholder needs. While both scenarios successfully accommodated ∼14% of EEZ (∼103,177 km^2^, 686.12 GW) for new OW projects, CZs in Scenario 1 included 1962.35 km^2^ of existing OWFs, 433 km^2^ of ports, and 1293 km^2^ of oil/gas fields compared to 119.91 km^2^, 3.7 km^2^, and 22 km^2^ in Scenario 2, indicating that greater spatial separation leads to achievable but slightly less ambitious conservation targets. The reduced overlap between conservation and anthropogenic activities in Scenario 2 can drive ecosystem recovery, especially for vulnerable habitats such as maërl beds (Steneck et al., 2002; Lotze et al., 2011; Burns et al., 2023). For instance, the spatial separation can result in large-scale recovery of seagrass habitats due to reduced nutrient pollution and eutrophication (Greening and Janicki, 2006; Reise and Kohlus, 2008). Consequently, Scenario 2 provides a balanced baseline for expanding the HPMA network beyond the current 985 km^2^, redesigning shipping routes, and accelerating progress towards national and global commitments (DEFRA, 2024).

Moreover, with a 74% overlap between the present MPA network and Scenario 2, redesigning the network requires only minor changes to zoning policy. The consistently higher proportion of undisturbed habitats in CZs than in MPAs and increased protection for coastal species complements the network without compromising renewable energy targets. While a large proportion of suitable habitat for macroalgae is undisturbed in offshore MPAs, over 60% remain open to fishing throughout the year, a gap more effectively addressed by Scenario 2 (Greenpeace UK, 2022).

Although implementing Scenario 2 may reduce the total amount of protected area and potentially result in several smaller reserves, evidence suggests that smaller, well-enforced MPAs can deliver large-scale ecological benefits (Rojo et al., 2019; Wilms et al., 2021).

Larger MPAs can reduce edge effects and enhance fish stock regeneration, but effective regulation across massive areas is often difficult (Walters, 2000; Roberts et al., 2001; De Santo, 2013). Adopting Scenario 2 as a baseline, upgrading protection statuses, expanding HPMAs, and reducing high-intensity overlaps could deliver greater long-term benefits by mimicking successful MPAs (see Edgar et al., (2014) and Burns et al., (2023)) and strategically distributing smaller sites as an initial compromise to reconcile competing priorities while building toward a more coherent network and meeting international commitments.

### Blue Carbon Potential

Our estimates suggest a range of 533,206.6-625,558.4 t C yr⁻¹ sequestered in 2026, which is nearly double earlier UK estimates (Burrows et al., 2024). Despite lower estimates for *Z. marina* and Phaeophyceae, the inclusion of subtidal habitats, modelled maërl habitat, and Rhodophyta results in substantially higher estimates (Burrows et al., 2024). While Burrows et al., (2024) used a fixed productivity rate (333.2 g C m^-2^ yr^-1^) for macroalgae, our estimates incorporated averages across all subtidal Phaeophyceae and quantified uncertainty. Similarly, our *Z. marina* estimates were based on UK-specific intertidal sequestration rates, while Burrows et al., (2024) used global averages, highlighting the need for region-specific data (do Amaral Camara Lima, 2020).

The high potential of maërl beds (88,152-92,035 t C yr⁻¹) reflects modelled suitable habitat rather than the currently mapped < 100 km^2^ of maërl beds (Simon-Nutbrown et al., 2020; Sheehy et al., 2024). Realising this potential is improbable without immediate protection of undisturbed areas and active restoration, given maërl’s slow growth rates and sensitivity to acidification, sedimentation, and climate change (Raven, 2018; Joshi and Farrell, 2020; Tuya et al., 2023). Even after excluding maërl beds, our inclusion of Rhodophyta yielded higher totals, representing the first UK-wide estimate for red macroalgal sequestration. Our undisturbed seagrass and Phaeophyceae estimates closely match mapped distributions (Yesson et al., 2015; Smale et al., 2016; Green et al., 2021; Burrows et al., 2024). However, further analysis is needed to determine whether this alignment reflects a methodological artefact, an upper bound on undisturbed habitat, or genuine model robustness.

Although the upper-limit sequestration from undisturbed habitats represents only ∼0.6% of annual UK emissions, this figure excludes wider ecosystem services, which can range from $87 – $147 km^-2^ yr^-1^ (Howard-Williams, 2022; Eger et al., 2023; Stamatiadou et al., 2025). Similarly, our restoration estimate of 110.9-122.3 t C yr⁻¹ by 2040 is likely conservative, as it omits these co-benefits and the economic viability of restoration, with benefit:cost ratios ranging from 1.1 -5 (Stewart-Sinclair et al., 2021; Carnell et al., 2025). While most projects to date have been small (median size: 0.04 ha), with site selection as a key determinant of success, our marine spatial plan and habitat maps provide a pathway to establish large-scale restoration or augmentation, provided technical barriers are overcome, which can result in conservation and economic benefits (Fraschetti et al., 2021; Eger et al., 2022; Danovaro et al., 2025). Therefore, we recommend protecting and augmenting existing undisturbed habitats while implementing adjacent restoration to increase connectivity and success rates.

The large uncertainties in our estimates, driven by limited data, indicate the need for improved measurements of marine carbon dynamics. For example, no UK-specific data exists for subtidal seagrass sequestration rates, with only one study addressing intertidal meadows (do Amaral Camara Lima, 2020). Future estimates can be refined through species-specific habitat models, growth curves, increased mapping, and studies on carbon dynamics.

However, given the high spatial variability in NPP and carbon burial, and the challenges of measurement and verification, we emphasise the adoption of uncertainty frameworks to define acceptable confidence thresholds and avoid the pitfall of perpetual monitoring.

### Limitations

Firstly, the difficulty in accounting for habitat overlaps between habitats and intertidal species and the coarse resolution of 500m, assuming full occupancy per pixel, may have inflated suitable habitat (Smale et al., 2016). Secondly, the absence of fine-scale substrate data likely overestimates suitable habitat extent, highlighting a need for higher-resolution benthic mapping. Third, sequestration estimates were limited to undisturbed areas, meaning our totals are likely conservative, while restoration scenarios assumed static habitat suitability rather than incorporating future suitability under different scenarios. Fourth, macroalgal sequestration rates were derived at the family level due to data scarcity. Lastly, additional features such as IUCN range maps or STAR data could improve our framework (Mair et al., 2021; IUCN, 2025).

### Policy Implications

Our marine spatial plan in Scenario 2 enables clear spatial separation between conflicting stakeholder interests with minor changes to the current MPA network. Leveraging this as a baseline for future policy is critical for aligning UK Net-Zero targets with competing stakeholder priorities. By altering the current network and achieving spatial separation of stakeholder use, marine habitats can recover without impacting other priorities. Our preliminary zoning plan provides a foundation that must be refined through stakeholder consultation to ensure comprehensive habitat coverage and to incorporate additional ecological layers, including seabirds, marine fauna, and benthic substrate.

The integration of OWFs within our plans illustrates how conservation and energy goals can be reconciled. With 41% of the European Union’s OW resource located in UK waters, the EEZ is critical to meeting the international target of 300 GW of OW capacity across the North Seas by 2050 (National Grid, 2024; The Crown Estate, 2024). Of the 22,428 km^2^ (149 GW) of technical potential identified in our ENZs in the North Sea, only a small fraction can be deployed (Archer and Jacobson, 2013; World Bank, 2019). However, based on UK’s North Sea pipeline (70 GW), developing 20% of this potential would enable the UK to contribute 33% of the 300 GW target (GEM Wiki, 2025). Leveraging the UK’s mature OW infrastructure, and advanced regulatory systems, alongside the declining costs of turbine installation/maintenance provides an opportunity to deploy high-capacity projects to accelerate global decarbonisation goals while positioning the UK as a central node in a pan-European renewable energy network (Elia et al., 2020; The Crown Estate, 2024; National Grid, 2024).

Moreover, given that this potential lies solely in the ENZ, energy goals can be achieved without impacting biodiversity, potentially enhancing biodiversity by leveraging synergies. Measures such as mandating ECOncrete in turbine foundations to create artificial reefs or promoting aquaculture co-location could ensure conservation across the broader seascape (Sella et al., 2021; WWF, 2025; Stockbridge et al., 2025). Furthermore, incorporating our findings within policy frameworks such as Marine Biodiversity Net Gain and the Offshore Wind Environmental Improvement Package, and aligning them with the EU Nature Market Roadmap and WWF’s Nature Positive Pathways, would ensure biodiversity is considered across the infrastructure lifecycle (CCC, 2022; DEFRA, 2023; DEFRA, 2025b; WWF, 2025; European Commission, 2025).

The spatial separation achieved in our plan is also aligned with the UK Marine Strategy, where only 2/15 Good Environmental Status (GES) targets have been met (DEFRA, 2025a). Our plan directly addresses 8/15 GES descriptors. For example, it mitigates the currently uncertain GES for underwater noise by reducing vessel density and oil/gas field and port overlaps, which can significantly disrupt marine biodiversity (Duarte et al., 2021; DEFRA, 2025a; Wilson et al., 2025). Implementing whole-site management strategies in the spatially isolated CZs, similar to Lyme Bay, could deliver wider ecosystem-wide benefits (Davies et al., 2022) through spillover effects (Da Silva et al., 2015), benefit fisheries (Kerwath et al., 2013), and help achieve GES targets (Solandt et al., 2020; Renn et al., 2024; DEFRA, 2025a).

Our blue carbon estimates, identification of priority habitats, and the accommodation of ∼37% of the EEZ as CZs highlight the high potential of NbS to accelerate progress towards policy commitments without undermining competing interests. In particular, our estimates support the inclusion of marine blue carbon ecosystems in national GHG inventories, where they are currently excluded or only partially accounted for (IPCC, 2014; CCC, 2022). Our methodology also provides a transferable framework for embedding natural capital into corporate decision-making via the Corporate Sustainability Reporting Directive (EP and CEU, 2022). This is critical in addressing NbS value mismatches and bridging climate finance gaps by engaging businesses to scale restoration through voluntary blue carbon markets (DEFRA, 2025c; Chausson et al., 2025).

Finally, our findings also have implications for broader policy innovation, including supporting a circular bioeconomy through sustainable macroalgal use (Lomartire et al., 2022). Protecting and restoring kelp habitats, identified as high-potential blue carbon sinks in our models, can synergistically reduce terrestrial emissions through dietary changes, kelp-based bioplastics, and bio-packaging, further encouraging marine conservation (Lomartire et al., 2022; De Souza Celente et al., 2023; Carr et al., 2025).

## Conclusion

This study presents the first integrated assessment of marine spatial planning, blue carbon potential, and restoration prioritisation across the UK EEZ. By coupling species-specific SDMs with spatial planning, we provide a preliminary framework for embedding marine NbS into Net-Zero policy. Despite conservative assumptions and data gaps, our findings show that resolving spatial conflicts and expanding the HPMA network can deliver measurable biodiversity gains, with only minor modifications to the current configuration. Top-down regulation, alignment with voluntary carbon markets, and stringent policy frameworks are essential to leverage these benefits, unlock climate finance, and help close the biodiversity funding gap while advancing global climate commitments.

