## Supplementary_Data for "Exploring marine habitat restoration opportunities and blue carbon sequestration potential in the United Kingdom"

**Table of Contents:**

| **Supplementary Methods** | Page 2 |
| --- | --- |
| **Supplementary Tables** | Page 6 |
| **Supplementary Figures** | Page 17 |
| **References** | Page 21 |

**Supplementary Methods**

*Species Distribution Models Algorithms*

MaxEnt uses presence-only data, generated background/pseudoabsence points, and covariates to output a probability distribution of suitable habitat (Phillips et al., 2006). MaxEnt fits functions using features derived from covariates, including linear, product, quadratic, hinge, threshold, and categorical classes (Elith et al., 2011). The linear feature class attempts to match mean values of covariates between presences and predicted presences, whereas the quadratic and product classes constrain the variance (Merow et al., 2013). The product feature class constrains the covariance of a predictor with other covariates, similar to the interaction term in regressions (Merow et al., 2013). Threshold and hinge features convert continuous predictors into binary predictors using a step and linear function, respectively (Phillips and Dudík, 2008). Categorical features convert categorical predictors into n binary features. Normalisation of coefficients is conducted to enable comparisons (Merow et al., 2013).

Boosted regression trees (BRT) combine decision trees and boosting algorithms (Elith et al., 2008). Boosting is a procedure employed to improve models through an iterative tree-fitting process based on sequentially optimising the loss function by incorporating weak learners to reduce residuals from poorly performing covariates from previous iterations (Schapire, 2003; Elith et al., 2008). While random forests and BRTs use decision trees, random forest trees are independently trained, unlike BRTs, where boosting allows for increased focus on fine-scale patterns in data (Breiman, 2001; Elith et al., 2008). BRTs require three parameters - tree complexity, bag fraction, and learning rate. The learning rate defines the contribution of each tree to the final model, with smaller values resulting in the evaluation of more trees, while the tree complexity refers to the number of splits per tree (Elith et al., 2008). The bag fraction specifies the proportion of training data used in each iteration and influences stochasticity in models.

*Marxan with Zones (MARZONE)*

MARZONE, an extension of Marxan, is a decision support tool for systematic conservation planning (Watts et al., 2009). MARZONE solves a ‘minimum set problem’ by assigning planning units (PU) to user-defined zones while meeting conservation targets at a minimum cost. Unlike Marxan, MARZONE allows the development of multiple zones with the option to assign zone-specific costs and targets, allowing practitioners to align the interests of multiple stakeholders. Consequently, MARZONE has been utilised in developing zoning plans for multiple ecosystems across different scales, ranging from developing plans for a 38 km^2^ marine reserve in Australia to 583,000 km^2^ in Borneo (Watts et al., 2009; Runting et al., 2015).

MARZONE aims to minimise the cost across PUs, the cost of boundaries between zones, and the penalties for not achieving targets (Equation S1). For instance, MARZONE can assign different costs for a conservation zones bordering fishing zones and partial conservation zones to ensure minimal boundaries between conservation and fishing zones.

$$Min\left( \sum_{PUs} Cost+\sum_{PUs} Boundary Costs+\sum_{PUs} (Boundary Length*Boundary Costs)+ \sum_{Features} (Feature Penalty Factor*Penalty) \right)$$

**Equation S1.** Objective function of Marxan with Zones. PUs refer to planning units

MARZONE optimises the objective function using a simulated annealing algorithm, which can be combined with an iterative improvement algorithm. he simulated annealing algorithm is an optimisation used to find a global optimum by stochastically accepting solutions that increase the objective function to prevent getting stuck in a local minimum (Henderson et al., 2003). The probability of accepting an increase in the objective function is influenced by a variable called temperature, which gradually decreases with each iteration until no increases in the objective function are accepted (Henderson et al., 2003). In MARZONE, this involves iteratively examining the change of adding/removing a random planning unit from the zone configuration (Serra-Sogas et al., 2020). In MARZONE, simulated annealing can be complemented by iterative improvement, which makes a random change to consider the impact on the objective function and continues until no change reduces the function (Serra-Sogas et al., 2020). The types of changes tested can include additions and removals (normal), reciprocal exchanges between planning units (swap), or paired additions and removals across all combinations (two-step), each offering increasing solution precision at the cost of computational complexity (Serra-Sogas et al., 2020).

*Marxan with Zones preparation and conditions*

MARZONE requires seven mandatory input files, with several optional files to include additional constraints. The EEZ was divided into 5 km x 5km square PUs followed by estimating the feature-specific overlap with each PU.

PU costs, derived using area as a proxy, were classified into six categories: Stakeholder, Oil, Energy, Conservation, Shipwreck, and NA Cost. Stakeholder costs incorporated ports and vessel routes, whereas oil costs included oil and gas fields and pipelines. In Scenario 1, Conservation costs were defined as the maximum value of MPAs or biodiversity features; in Scenario 2, they were limited to biodiversity values. Shipwreck costs were treated independently, reflecting their relevance solely to the ENZ. The NA cost was set at 10 for units lacking features and 0 for those with features, to ensure that featureless units were allocated to either CZ or OPZ. All costs, except the NA cost, were normalised to the range [0,1].

Zone-specific cost multipliers are detailed in Table S3. Shipwreck costs were applied only to the ENZ assuming increased engineering and logistical complexities. To ensure inclusion of relevant features to our defined zones, we included zone contributions to feature targets (Table S4). CZs contributed 100% toward biodiversity targets, while Open, Oil, and Energy zones contributed 30%, 0%, and 20%, respectively, based on assessed compatibilities (Werner et al., 2024). In Scenario 1, 10,450 PUs were locked into the CZ (Equation S2).

We defined zone boundary costs to increase spatial clumping of zones (Table S5).

$$IF(AND(MPA > 12.5, IFERROR(MPA / (Oil + MPA + Wind\_Farm), 0) \geq0.5), 1, 0)$$

**Equation S2.** Microsoft Excel formula used to identify planning units (25 km^2^) locked into the conservation zone under Scenario 1. A unit was locked if Marine Protected Area (MPA) coverage exceeded 12.5 km^2^ and MPAs comprised ≥50% of the combined MPA, oil, and wind farm area within the unit.

We defined zone boundary costs to increase spatial clumping of zones (Table S5). Costs were set at -2 within zones to encourage clumping, while inter-zone costs were set to discourage boundaries between non-synergistic zones.

Several key parameters, including boundary costs, feature penalty factors, and zone-specific cost multipliers iteratively calibrated from prior runs to ensure balanced trade-offs between ecological representation, spatial efficiency, and stakeholder priorities.

MARZONE was executed using the simulated annealing algorithm followed by iterative improvement for 1000 iterations, constrained by computational limitations.

**Table S1.** Fitted variogram model parameters used in universal kriging of log-transformed seabed flow velocity with depth as a covariate. The model consists of a nugget effect and a Matérn (“Ste”) model. Psill – partial sill, ang -angle, anis – anisotropy.

| **model** | **psill** | **range** | **kappa** | **ang1** | **ang2** | **ang3** | **anis1** | **anis2** |
| --- | --- | --- | --- | --- | --- | --- | --- | --- |
| Nug | 0 | 0 | 0 | 0 | 0 | 0 | 1 | 1 |
| Ste | 0.294597 | 198809.5 | 0.4 | 0 | 0 | 0 | 1 | 1 |

**Table S2.** Environmental thresholds used to map habitat suitability for brown and red macroalgae (Duarte et al., 2022)

| **Variable** | **Phaeophyceae** | **Rhodophyta** |
| --- | --- | --- |
| Seabed Temperature (C°) | <28 | <35 |
| Seabed Salinity (PSU) | >7 | >7 |
| Permanent Ice Thickness (m) | 0 | 0 |
| Light at Bottom (moles photons m^-2^day^-1^) | 0.0277 | 0.0001 |
| Nitrate at bottom (mmol/m^3^) | 0.002- 94.820 | 0.002- 94.820 |

**Table S3.** Costs and multipliers applied to each zone. Normalised areas of features were used as proxy for costs. If no feature was present in a planning unit, a cost of 10 was applied (NA Cost). Stakeholder Costs - Vessel Density and Ports, Conservation Costs – Maximum of all biodiversity features and MPAs

| **Zone** | **Cost** | **Multiplier** |
| --- | --- | --- |
| Open | Oil & Pipeline | 0.2 |
| Open | Energy | 0.2 |
| Open | Conservation | 0.2 |
| Open | NA Cost | 9 |
| Conservation | Stakeholder | 2 |
| Conservation | Oil & Pipeline | 10 |
| Conservation | Energy | 5 |
| Conservation | Conservation | 0.05 |
| Conservation | NA Cost | 8.9 |
| Oil | Stakeholder | 2 |
| Oil | Oil & Pipeline | 0.05 |
| Oil | Energy | 2 |
| Oil | Conservation | 2 |
| Oil | NA Cost | 10 |
| Energy | Stakeholder | 1 |
| Energy | Energy | 0.05 |
| Energy | Conservation | 2 |
| Energy | Shipwrecks | 2 |
| Energy | NA Cost | 10 |

**Table S4.** Contribution of each zone to feature targets in Scenarios 1 and 2

| Zone | Feature | Scenario 1 | Scenario 2 |
| --- | --- | --- | --- |
| Open | *Zostera marina* | 0.3 | 0.3 |
| Open | Rhodophyta | 0.3 | 0.3 |
| Open | Phaeophycae | 0.3 | 0.3 |
| Open | *Phymatolithon calcaereum* | 0.3 | 0.3 |
| Open | *Lithothamnion corallioides* | 0.3 | 0.3 |
| Open | *Lithothamnion glaciale* | 0.3 | 0.3 |
| Open | Offshore Wind Potential | 0.15 | 0.15 |
| Open | Wind Farms | 0.15 | 0.15 |
| Open | Oil Fields | 0.15 | 0.15 |
| Open | Pipelines | 1 | 1 |
| Open | Ports | 1 | 1 |
| Open | Vessels | 1 | 1 |
| Open | Marine Protected Areas | 1 | NA |
| Open | Shipwrecks | 0 | 1 |
| Open | Protected Areas | NA | 0 |
| Conservation | *Zostera marina* | 1 | 1 |
| Conservation | Rhodophyta | 1 | 1 |
| Conservation | Phaeophycae | 1 | 1 |
| Conservation | *Phymatolithon calcaereum* | 1 | 1 |
| Conservation | *Lithothamnion corallioides* | 1 | 1 |
| Conservation | *Lithothamnion glaciale* | 1 | 1 |
| Conservation | Offshore Wind Potential | 0.3 | 0.3 |
| Conservation | Wind Farms | 0.3 | 0.3 |
| Conservation | Oil Fields | 0 | 0 |
| Conservation | Pipelines | 1 | 1 |
| Conservation | Ports | 0 | 0 |
| Conservation | Vessels | 1 | 1 |
| Conservation | Marine Protected Areas | 1 | 1 |
| Conservation | Shipwrecks | 1 | NA |
| Conservation | Protected Areas | NA | 1 |
| Oil | *Zostera marina* | 0 | 0 |
| Oil | Rhodophyta | 0 | 0 |
| Oil | Phaeophycae | 0 | 0 |
| Oil | *Phymatolithon calcaereum* | 0 | 0 |
| Oil | *Lithothamnion corallioides* | 0 | 0 |
| Oil | *Lithothamnion glaciale* | 0 | 0 |
| Oil | Offshore Wind Potential | 1 | 1 |
| Oil | Wind Farms | 1 | 1 |
| Oil | Oil Fields | 1 | 1 |
| Oil | Pipelines | 1 | 1 |
| Oil | Ports | 1 | 1 |
| Oil | Vessels | 1 | 1 |
| Oil | Marine Protected Areas | 0 | NA |
| Oil | Shipwrecks | 1 | 1 |
| Oil | Protected Areas | NA | 0 |
| Energy | *Zostera marina* | 0.2 | 0.2 |
| Energy | Rhodophyta | 0.2 | 0.2 |
| Energy | Phaeophycae | 0.2 | 0.2 |
| Energy | *Phymatolithon calcaereum* | 0.2 | 0.2 |
| Energy | *Lithothamnion corallioides* | 0.2 | 0.2 |
| Energy | *Lithothamnion glaciale* | 0.2 | 0.2 |
| Energy | Offshore Wind Potential | 1 | 1 |
| Energy | Wind Farms | 1 | 1 |
| Energy | Oil Fields | 1 | 1 |
| Energy | Pipelines | 1 | 1 |
| Energy | Ports | 1 | 1 |
| Energy | Vessels | 1 | 1 |
| Energy | Marine Protected Areas | 1 | NA |
| Energy | Shipwrecks | 0 | 0 |
| Energy | Protected Areas | NA | 0 |

**Table S5.** Boundary costs across zones across Scenarios. Lower costs encourage spatial clumping

| **Zone 1** | **Zone 2** | **Boundary Cost** |
| --- | --- | --- |
| Open | Open | -2 |
| Open | Conservation | 20 |
| Open | Oil | 0 |
| Open | Energy | 0 |
| Conservation | Conservation | -5 |
| Conservation | Oil | 10 |
| Conservation | Energy | 6 |
| Oil | Oil | -2 |
| Oil | Energy | 2 |
| Energy | Energy | -2 |

**Table S6a.** Feature representation summary from the best solution under Scenario 1 showing target values and held/contributing amounts across each planning MPM indicates the Minimum Proportion Met, with targets considered met if ≥97% of the total target was achieved. Contributing amounts refer to the portion of each feature within the zone that actively contributes toward the target.

| Feature Name | Target | Amount Held Open | Contributing Amount Held Open | Target Conservation | Amount Held Conservation | Contributing Amount Held Conservation | Target Oil | Amount Held Oil | Contributing Amount Held Oil | Target Energy | Amount Held Energy | Contributing Amount Held Energy | MPM |
| --- | --- | --- | --- | --- | --- | --- | --- | --- | --- | --- | --- | --- | --- |
| Shipwrecks | 3673.811 | 1995.624 | 1995.624 | 0 | 1569.526 | 1569.526 | 0 | 108.6468 | 108.6468 | 0 | 0.0141 | 0 | 1.000 |
| MPAs | 302061.9 | 3582.166 | 3582.166 | 241649.5 | 257570.1 | 257570.1 | 0 | 20515.65 | 0 | 0 | 20393.96 | 20393.96 | 0.932 |
| Vessels | 81602.27 | 29121.25 | 29121.25 | 0 | 21117.93 | 21117.93 | 0 | 17532.57 | 17532.57 | 0 | 13830.52 | 13830.52 | 1.000 |
| Ports | 1676.308 | 1103.617 | 1103.617 | 0 | 433.9362 | 0 | 0 | 81.91291 | 81.91291 | 0 | 56.8419 | 56.8419 | 0.741 |
| Pipelines | 91411.14 | 13948.78 | 13948.78 | 0 | 7453.021 | 7453.021 | 0 | 46430.19 | 46430.19 | 0 | 23579.14 | 23579.14 | 1.000 |
| Oil | 89388.93 | 356.2165 | 53.43247 | 0 | 1293.24 | 0 | 71511.14 | 71510.39 | 71510.39 | 0 | 16229.08 | 16229.08 | 0.982 |
| Wind Farms | 68143.84 | 444.8827 | 66.73241 | 0 | 1962.36 | 588.7079 | 0 | 11222.54 | 11222.54 | 54515.07 | 54514.05 | 54514.05 | 0.974 |
| Wind Potential | 104900.5 | 114040.8 | 17106.13 | 0 | 16.33019 | 4.899056 | 0 | 2289.468 | 2289.468 | 103176.3 | 103177.4 | 103177.4 | 1.000 |
| Glaciale | 865.1622 | 1637.876 | 491.3629 | 432.5811 | 989.7987 | 989.7987 | 0 | 37.09253 | 0 | 0 | 219.1066 | 43.82131 | 1.000 |
| Coralloides | 120.1289 | 287.3305 | 86.19914 | 60.06447 | 86.67847 | 86.67847 | 0 | 2.603544 | 0 | 0 | 23.81735 | 4.763469 | 1.000 |
| Calcaereum | 1234.945 | 2470.614 | 741.1842 | 617.4726 | 1117.596 | 1117.596 | 0 | 74.73281 | 0 | 0 | 453.5413 | 90.70825 | 1.000 |
| Phaeophycae | 24208.73 | 26740.61 | 8022.182 | 12104.37 | 28037.85 | 28037.85 | 0 | 9554.982 | 0 | 0 | 16362.34 | 3272.468 | 1.000 |
| Rhodophyta | 102657.8 | 140041.6 | 42012.48 | 51328.92 | 89006.35 | 89006.35 | 0 | 50087.63 | 0 | 0 | 63057.27 | 12611.45 | 1.000 |
| Zostera | 836.5225 | 1727.036 | 518.1107 | 418.2612 | 818.7759 | 818.7759 | 0 | 90.39409 | 0 | 0 | 152.2026 | 30.44052 | 1.000 |

**Table S6b.** Feature representation summary from the best solution under Scenario 2 showing target values and held/contributing amounts across each planning MPM indicates the Minimum Proportion Met, with targets considered met if ≥98% of the total target was achieved. Contributing amounts refer to the portion of each feature within the zone that actively contributes toward the target.

| Feature Name | Target | Amount Held Open | Contributing Amount Held Open | Target Conservation | Amount Held Conservation | Contributing Amount Held Conservation | Target Oil | Amount Held Oil | Contributing Amount Held Oil | Target Energy | Amount Held Energy | Contributing Amount Held Energy | MPM |
| --- | --- | --- | --- | --- | --- | --- | --- | --- | --- | --- | --- | --- | --- |
| Protected Area | 218170.3 | 178745.5 | 0 | 218167.3 | 269342.9 | 269342.9 | 0 | 89304.64 | 0 | 0 | 189831.3 | 0 | 1.000 |
| Shipwrecks | 3673.811 | 2186.083 | 2186.083 | 0 | 1381.875 | 1381.875 | 0 | 89.41758 | 89.41758 | 0 | 16.43591 | 0 | 0.996 |
| Vessels | 81602.27 | 41869.9 | 41869.9 | 0 | 7733.325 | 7733.325 | 0 | 17054.68 | 17054.68 | 0 | 14944.36 | 14944.36 | 1.000 |
| Ports | 1676.308 | 1507.413 | 1507.413 | 0 | 3.737088 | 0 | 0 | 29.50755 | 29.50755 | 0 | 135.6504 | 135.6504 | 0.998 |
| Pipelines | 91411.14 | 19299.69 | 19299.69 | 0 | 101.9735 | 101.9735 | 0 | 46600.03 | 46600.03 | 0 | 25409.45 | 25409.45 | 1.000 |
| Oil | 89388.93 | 353.8629 | 53.07944 | 0 | 22.24207 | 0 | 71511.14 | 71508.18 | 71508.18 | 0 | 17504.65 | 17504.65 | 0.996 |
| Wind Farms | 68143.84 | 1007.01 | 151.0515 | 0 | 119.9191 | 35.97573 | 0 | 12503.09 | 12503.09 | 54515.07 | 54513.82 | 54513.82 | 0.986 |
| Wind Potential | 104900.5 | 113879.9 | 17081.98 | 0 | 710.2616 | 213.0785 | 0 | 1756.077 | 1756.077 | 103176.3 | 103177.8 | 103177.8 | 1.000 |
| Glaciale | 865.1622 | 1933.107 | 579.9322 | 576.7748 | 694.7095 | 694.7095 | 0 | 18.87425 | 0 | 0 | 237.1832 | 47.43664 | 1.000 |
| Coralloides | 120.1289 | 272.635 | 81.7905 | 80.08597 | 100.212 | 100.212 | 0 | 1.079475 | 0 | 0 | 26.5033 | 5.30066 | 1.000 |
| Calcaereum | 1234.945 | 2778.066 | 833.4197 | 823.2968 | 839.12 | 839.12 | 0 | 58.72245 | 0 | 0 | 440.576 | 88.1152 | 1.000 |
| Phaeophycae | 24208.73 | 34390.98 | 10317.29 | 16139.16 | 19060.19 | 19060.19 | 0 | 10380.13 | 0 | 0 | 16864.48 | 3372.896 | 1.000 |
| Rhodophyta | 102657.8 | 158851.4 | 47655.43 | 68438.57 | 68438.61 | 68438.61 | 0 | 52059.42 | 0 | 0 | 62843.37 | 12568.67 | 1.000 |
| Zostera | 836.5225 | 1823.166 | 546.9498 | 557.6816 | 725.7288 | 725.7288 | 0 | 63.28964 | 0 | 0 | 176.2239 | 35.24479 | 1.000 |

**Table S7.** Selection frequency metrics for planning units (PUs) across zones under both Scenario 1 (MPA) and 2 (Free). Irreplaceable PUs are defined as those selected in 100% of runs, while high-priority units are those selected in at least 90% of runs.

| **Zone** | **Scenario** | **Total Selected** | **Irreplaceable PUs** | **% PUs Irreplaceable** | **High Priority PUs** | **% PUs high priority** |
| --- | --- | --- | --- | --- | --- | --- |
| Conservation | Free | 14109 | 10838 | 76.82 | 10850 | 76.9 |
| Conservation | MPA | 12628 | 12600 | 99.78 | 12600 | 99.78 |
| Energy | Free | 16483 | 4487 | 27.22 | 4598 | 27.9 |
| Energy | MPA | 16308 | 4377 | 26.84 | 4508 | 27.64 |
| Oil | Free | 6471 | 16 | 0.25 | 2416 | 37.34 |
| Oil | MPA | 6378 | 10 | 0.16 | 3336 | 52.3 |
| Open | Free | 10335 | 1089 | 10.54 | 6238 | 60.36 |
| Open | MPA | 8742 | 2247 | 25.7 | 7347 | 84.04 |

**Table S8.** Spearman’s rank correlation between selection frequency across Scenarios. Strong positive correlations (ρ > 0.89) were observed in all zones (p < 0.001)

| **Zone** | **Spearman_rho** | **p_value** |
| --- | --- | --- |
| Open | 0.874 | <0.001 |
| Conservation | 0.906 | <0.001 |
| Oil | 0.958 | <0.001 |
| Energy | 0.963 | <0.001 |

**Table S9.** Suitable habitat in Undisturbed regions within the Conservation Zone. Area in MPA includes undisturbed habitat in the Conservation Zone overlapping with MPAs. MPA- Marine Protected Area. All areas are in km^2^

| **Scenario** | **Feature** | **Suitable habitat in conservation zone** | **Area undisturbed** | **% undisturbed** | **Undisturbed Area in MPA** | **% in MPA** |
| --- | --- | --- | --- | --- | --- | --- |
| Scenario 1 | Calcareum | 1117.593 | 139.323 | 12.47 | 76.291 | 6.83 |
| Scenario 2 | Calcareum | 915.852 | 144.055 | 15.73 | 75.858 | 8.28 |
| Scenario 1 | Corallioides | 86.683 | 9.039 | 10.43 | 2.27 | 2.62 |
| Scenario 2 | Corallioides | 104.739 | 9.039 | 8.63 | 2.27 | 2.17 |
| Scenario 1 | Glaciale | 989.797 | 120.391 | 12.16 | 80.104 | 8.09 |
| Scenario 2 | Glaciale | 772.5 | 121.158 | 15.68 | 79.922 | 10.35 |
| Scenario 1 | Phaeophyceae | 28037.85 | 8830.805 | 31.5 | 7609.041 | 27.14 |
| Scenario 2 | Phaeophyceae | 18797.33 | 8572.932 | 45.61 | 7094.984 | 37.74 |
| Scenario 1 | Rhodophyta | 89006.35 | 47632.25 | 53.52 | 39453.701 | 44.33 |
| Scenario 2 | Rhodophyta | 68438.57 | 46050.71 | 67.29 | 35983.269 | 52.58 |
| Scenario 1 | Zostera | 818.776 | 149.868 | 18.3 | 53.652 | 6.55 |
| Scenario 2 | Zostera | 740.008 | 150.579 | 20.35 | 53.652 | 7.25 |

**Table S10.** Blue carbon sequestration potential (t C yr⁻¹) from 2026–2040 for each scenario, based on Monte Carlo simulations (n = 1000). CI refers to 95% confidence intervals and IQR refers to interquartile Range

| Species | Scenario | Mean | Lower CI | Upper CI | SD | Median | IQR | Lower CI | Upper CI | Method |
| --- | --- | --- | --- | --- | --- | --- | --- | --- | --- | --- |
| Calcaereum | 2 | 47674.00 | 47367.56 | 47980.52 | 4938.90 | 47706.20 | 6994.00 | 47350.31 | 48297.73 | Normal CI |
| Calcaereum | 1 | 46108.10 | 45811.64 | 46404.47 | 4776.60 | 46139.10 | 6764.30 | 45794.97 | 46711.26 | Normal CI |
| Corallioides | 2 | 2975.60 | 2957.18 | 2993.99 | 296.50 | 2982.10 | 399.70 | 2957.23 | 2999.88 | Normal CI |
| Corallioides | 1 | 2975.60 | 2957.18 | 2993.99 | 296.50 | 2982.10 | 399.70 | 2957.23 | 2999.88 | Normal CI |
| Glaciale | 2 | 39868.00 | 39629.98 | 40106.09 | 3836.20 | 39839.50 | 4914.80 | 39652.79 | 40109.00 | Normal CI |
| Glaciale | 1 | 39615.70 | 39379.11 | 39852.20 | 3811.90 | 39587.30 | 4883.70 | 39401.77 | 39855.09 | Normal CI |
| Phaeophycae | 2 | 272604.90 | 253539.07 | 291913.92 | 314265.60 | 165424.30 | 301397.20 | 148266.43 | 181111.22 | Bootstrap CI |
| Phaeophycae | 1 | 280804.70 | 260844.73 | 301053.68 | 323718.70 | 170400.20 | 310463.20 | 152726.27 | 186559.03 | Bootstrap CI |
| Rhodophyta | 2 | 385332.20 | 353919.28 | 418351.07 | 513991.30 | 184219.40 | 446635.60 | 168620.32 | 209657.33 | Bootstrap CI |
| Rhodophyta | 1 | 398941.60 | 364221.99 | 430918.06 | 532144.80 | 190725.80 | 462410.20 | 174575.78 | 217062.16 | Bootstrap CI |
| Zostera Marina | 2 | 10159.60 | 9866.92 | 10452.25 | 4716.30 | 10074.30 | 6532.40 | 9602.59 | 10326.28 | Normal CI |
| Zostera Marina | 1 | 10111.60 | 9820.34 | 10402.91 | 4694.00 | 10026.80 | 6501.60 | 9557.24 | 10277.52 | Normal CI |
| Calcaereum | 2 | 47675.20 | 47368.76 | 47981.74 | 4939.00 | 47707.80 | 6995.20 | 47351.67 | 48299.55 | Normal CI |
| Calcaereum | 1 | 46109.30 | 45812.84 | 46405.68 | 4776.80 | 46140.70 | 6765.40 | 45796.33 | 46713.08 | Normal CI |
| Corallioides | 2 | 2976.80 | 2958.37 | 2995.19 | 296.60 | 2983.00 | 400.00 | 2958.72 | 3000.86 | Normal CI |
| Corallioides | 1 | 2976.80 | 2958.37 | 2995.19 | 296.60 | 2983.00 | 400.00 | 2958.72 | 3000.86 | Normal CI |
| Glaciale | 2 | 39869.20 | 39631.16 | 40107.28 | 3836.30 | 39840.60 | 4914.20 | 39654.04 | 40109.83 | Normal CI |
| Glaciale | 1 | 39616.80 | 39380.29 | 39853.39 | 3812.00 | 39588.40 | 4883.10 | 39403.02 | 39855.92 | Normal CI |
| Phaeophycae | 2 | 272610.70 | 251592.73 | 292561.27 | 314290.40 | 165424.30 | 301397.20 | 148266.43 | 181111.66 | Bootstrap CI |
| Phaeophycae | 1 | 280810.60 | 259561.53 | 300596.33 | 323743.40 | 170400.30 | 310463.20 | 152726.28 | 186559.47 | Bootstrap CI |
| Rhodophyta | 2 | 385333.00 | 352452.24 | 418012.42 | 513992.00 | 184219.40 | 446635.60 | 168620.39 | 209657.36 | Bootstrap CI |
| Rhodophyta | 1 | 398942.50 | 365811.18 | 430731.73 | 532145.50 | 190725.80 | 462410.20 | 174575.84 | 217062.19 | Bootstrap CI |
| Zostera Marina | 2 | 10161.80 | 9868.98 | 10454.68 | 4719.20 | 10074.50 | 6532.10 | 9604.01 | 10326.28 | Normal CI |
| Zostera Marina | 1 | 10113.90 | 9822.40 | 10405.33 | 4696.90 | 10027.00 | 6501.20 | 9558.67 | 10277.52 | Normal CI |
| Calcaereum | 2 | 47676.50 | 47370.03 | 47983.02 | 4939.10 | 47708.60 | 6995.10 | 47352.70 | 48300.66 | Normal CI |
| Calcaereum | 1 | 46110.50 | 45814.11 | 46406.96 | 4776.90 | 46141.60 | 6765.40 | 45797.37 | 46714.19 | Normal CI |
| Corallioides | 2 | 2978.00 | 2959.62 | 2996.45 | 296.80 | 2984.70 | 401.30 | 2959.68 | 3002.03 | Normal CI |
| Corallioides | 1 | 2978.00 | 2959.62 | 2996.45 | 296.80 | 2984.70 | 401.30 | 2959.68 | 3002.03 | Normal CI |
| Glaciale | 2 | 39870.40 | 39632.37 | 40108.51 | 3836.40 | 39842.30 | 4914.50 | 39654.94 | 40111.37 | Normal CI |
| Glaciale | 1 | 39618.10 | 39381.50 | 39854.62 | 3812.10 | 39590.10 | 4883.40 | 39403.92 | 39857.46 | Normal CI |
| Phaeophycae | 2 | 272614.30 | 252065.37 | 291622.98 | 314294.10 | 165427.10 | 301397.20 | 148266.43 | 181111.66 | Bootstrap CI |
| Phaeophycae | 1 | 280814.20 | 260746.31 | 299240.31 | 323747.10 | 170403.10 | 310463.20 | 152726.28 | 186559.47 | Bootstrap CI |
| Rhodophyta | 2 | 385334.00 | 354096.32 | 417055.00 | 513993.20 | 184219.40 | 446635.60 | 168620.40 | 209657.36 | Bootstrap CI |
| Rhodophyta | 1 | 398943.40 | 362160.74 | 430864.38 | 532146.70 | 190725.80 | 462410.20 | 174575.85 | 217062.19 | Bootstrap CI |
| Zostera Marina | 2 | 10164.30 | 9871.33 | 10457.23 | 4720.90 | 10074.50 | 6523.00 | 9604.08 | 10328.67 | Normal CI |
| Zostera Marina | 1 | 10116.30 | 9824.74 | 10407.88 | 4698.60 | 10027.00 | 6492.20 | 9558.74 | 10279.91 | Normal CI |
| Calcaereum | 2 | 47677.80 | 47371.33 | 47984.34 | 4939.20 | 47709.80 | 6995.40 | 47353.89 | 48302.30 | Normal CI |
| Calcaereum | 1 | 46111.80 | 45815.41 | 46408.28 | 4777.00 | 46142.80 | 6765.70 | 45798.56 | 46715.84 | Normal CI |
| Corallioides | 2 | 2979.30 | 2960.92 | 2997.77 | 296.90 | 2985.50 | 401.90 | 2960.50 | 3003.77 | Normal CI |
| Corallioides | 1 | 2979.30 | 2960.92 | 2997.77 | 296.90 | 2985.50 | 401.90 | 2960.50 | 3003.77 | Normal CI |
| Glaciale | 2 | 39871.70 | 39633.66 | 40109.81 | 3836.50 | 39844.00 | 4914.20 | 39657.11 | 40113.29 | Normal CI |
| Glaciale | 1 | 39619.40 | 39382.79 | 39855.92 | 3812.20 | 39591.80 | 4883.10 | 39406.09 | 39859.38 | Normal CI |
| Phaeophycae | 2 | 272618.20 | 253020.33 | 292954.61 | 314294.80 | 165427.20 | 301397.90 | 148266.65 | 181111.72 | Bootstrap CI |
| Phaeophycae | 1 | 280818.10 | 260831.24 | 300776.86 | 323747.80 | 170403.10 | 310463.90 | 152726.49 | 186559.53 | Bootstrap CI |
| Rhodophyta | 2 | 385335.30 | 348580.30 | 418777.98 | 513994.20 | 184219.40 | 446690.90 | 168620.40 | 209657.36 | Bootstrap CI |
| Rhodophyta | 1 | 398944.70 | 363705.88 | 434652.87 | 532147.70 | 190725.80 | 462466.60 | 174575.85 | 217062.19 | Bootstrap CI |
| Zostera Marina | 2 | 10165.60 | 9872.58 | 10458.53 | 4721.30 | 10075.00 | 6523.00 | 9608.49 | 10328.67 | Normal CI |
| Zostera Marina | 1 | 10117.60 | 9826.00 | 10409.19 | 4699.00 | 10027.40 | 6492.20 | 9563.12 | 10279.91 | Normal CI |
| Calcaereum | 2 | 47679.20 | 47372.72 | 47985.75 | 4939.40 | 47711.00 | 6995.20 | 47355.68 | 48304.56 | Normal CI |
| Calcaereum | 1 | 46113.20 | 45816.80 | 46409.69 | 4777.10 | 46143.90 | 6765.50 | 45800.35 | 46718.09 | Normal CI |
| Corallioides | 2 | 2980.70 | 2962.29 | 2999.15 | 297.10 | 2986.80 | 401.60 | 2961.91 | 3005.06 | Normal CI |
| Corallioides | 1 | 2980.70 | 2962.29 | 2999.15 | 297.10 | 2986.80 | 401.60 | 2961.91 | 3005.06 | Normal CI |
| Glaciale | 2 | 39873.10 | 39635.05 | 40111.21 | 3836.70 | 39845.80 | 4914.50 | 39658.33 | 40114.61 | Normal CI |
| Glaciale | 1 | 39620.70 | 39384.17 | 39857.32 | 3812.40 | 39593.60 | 4883.40 | 39407.31 | 39860.70 | Normal CI |
| Phaeophycae | 2 | 272627.40 | 252506.84 | 292006.00 | 314306.00 | 165427.20 | 301398.00 | 148266.67 | 181111.73 | Bootstrap CI |
| Phaeophycae | 1 | 280827.30 | 261561.22 | 301756.99 | 323759.10 | 170403.20 | 310464.00 | 152726.51 | 186559.54 | Bootstrap CI |
| Rhodophyta | 2 | 385337.30 | 351550.91 | 416698.46 | 513998.00 | 184219.50 | 446690.90 | 168620.40 | 209657.36 | Bootstrap CI |
| Rhodophyta | 1 | 398946.70 | 365592.15 | 431146.00 | 532151.50 | 190725.90 | 462466.60 | 174575.85 | 217062.19 | Bootstrap CI |
| Zostera Marina | 2 | 10168.20 | 9875.15 | 10461.24 | 4722.40 | 10080.80 | 6523.00 | 9608.49 | 10343.95 | Normal CI |
| Zostera Marina | 1 | 10120.20 | 9828.56 | 10411.89 | 4700.10 | 10033.20 | 6492.20 | 9563.13 | 10295.11 | Normal CI |
| Calcaereum | 2 | 47680.70 | 47374.18 | 47987.23 | 4939.50 | 47712.50 | 6996.60 | 47357.52 | 48306.31 | Normal CI |
| Calcaereum | 1 | 46114.70 | 45818.26 | 46411.17 | 4777.30 | 46145.40 | 6766.90 | 45802.19 | 46719.84 | Normal CI |
| Corallioides | 2 | 2982.20 | 2963.77 | 3000.65 | 297.20 | 2988.30 | 401.30 | 2962.86 | 3006.54 | Normal CI |
| Corallioides | 1 | 2982.20 | 2963.77 | 3000.65 | 297.20 | 2988.30 | 401.30 | 2962.86 | 3006.54 | Normal CI |
| Glaciale | 2 | 39874.60 | 39636.49 | 40112.67 | 3836.80 | 39847.20 | 4914.40 | 39659.74 | 40116.39 | Normal CI |
| Glaciale | 1 | 39622.20 | 39385.61 | 39858.78 | 3812.50 | 39595.00 | 4883.30 | 39408.72 | 39862.48 | Normal CI |
| Phaeophycae | 2 | 272633.00 | 253374.01 | 292075.69 | 314308.70 | 165427.20 | 301397.80 | 148266.68 | 181111.73 | Bootstrap CI |
| Phaeophycae | 1 | 280832.90 | 259666.49 | 301287.84 | 323761.70 | 170403.20 | 310463.80 | 152726.52 | 186559.54 | Bootstrap CI |
| Rhodophyta | 2 | 385339.90 | 352752.89 | 416863.05 | 514004.40 | 184219.50 | 446691.00 | 168620.92 | 209657.36 | Bootstrap CI |
| Rhodophyta | 1 | 398949.30 | 363958.87 | 431256.91 | 532157.90 | 190725.90 | 462466.70 | 174576.38 | 217062.19 | Bootstrap CI |
| Zostera Marina | 2 | 10172.30 | 9879.00 | 10465.53 | 4725.90 | 10090.30 | 6523.50 | 9633.51 | 10345.17 | Normal CI |
| Zostera Marina | 1 | 10124.30 | 9832.42 | 10416.19 | 4703.70 | 10042.70 | 6492.70 | 9588.17 | 10296.32 | Normal CI |
| Calcaereum | 2 | 47682.30 | 47375.74 | 47988.80 | 4939.70 | 47713.90 | 6997.70 | 47358.95 | 48307.01 | Normal CI |
| Calcaereum | 1 | 46116.30 | 45819.82 | 46412.75 | 4777.50 | 46146.90 | 6767.90 | 45803.62 | 46720.55 | Normal CI |
| Corallioides | 2 | 2983.70 | 2965.28 | 3002.18 | 297.30 | 2990.00 | 402.00 | 2964.57 | 3008.46 | Normal CI |
| Corallioides | 1 | 2983.70 | 2965.28 | 3002.18 | 297.30 | 2990.00 | 402.00 | 2964.57 | 3008.46 | Normal CI |
| Glaciale | 2 | 39876.10 | 39638.03 | 40114.23 | 3836.90 | 39847.80 | 4915.60 | 39660.71 | 40117.03 | Normal CI |
| Glaciale | 1 | 39623.80 | 39387.16 | 39860.34 | 3812.70 | 39595.60 | 4884.50 | 39409.69 | 39863.13 | Normal CI |
| Phaeophycae | 2 | 272639.40 | 253673.48 | 291265.10 | 314312.90 | 165427.20 | 301398.70 | 148266.72 | 181111.75 | Bootstrap CI |
| Phaeophycae | 1 | 280839.30 | 259593.75 | 301245.45 | 323765.90 | 170403.20 | 310464.70 | 152726.56 | 186559.56 | Bootstrap CI |
| Rhodophyta | 2 | 385344.00 | 354247.30 | 416882.58 | 514014.30 | 184221.40 | 446691.00 | 168621.39 | 209657.43 | Bootstrap CI |
| Rhodophyta | 1 | 398953.40 | 364252.49 | 431816.58 | 532167.80 | 190727.80 | 462466.70 | 174576.84 | 217062.26 | Bootstrap CI |
| Zostera Marina | 2 | 10177.80 | 9884.46 | 10471.23 | 4727.90 | 10096.80 | 6539.40 | 9636.49 | 10345.19 | Normal CI |
| Zostera Marina | 1 | 10129.90 | 9837.87 | 10421.88 | 4705.60 | 10049.10 | 6508.50 | 9591.14 | 10296.34 | Normal CI |
| Calcaereum | 2 | 47683.90 | 47377.38 | 47990.46 | 4939.90 | 47715.90 | 6997.60 | 47359.50 | 48308.39 | Normal CI |
| Calcaereum | 1 | 46117.90 | 45821.46 | 46414.41 | 4777.60 | 46148.90 | 6767.90 | 45804.17 | 46721.92 | Normal CI |
| Corallioides | 2 | 2985.30 | 2966.88 | 3003.81 | 297.50 | 2991.60 | 401.70 | 2966.66 | 3009.88 | Normal CI |
| Corallioides | 1 | 2985.30 | 2966.88 | 3003.81 | 297.50 | 2991.60 | 401.70 | 2966.66 | 3009.88 | Normal CI |
| Glaciale | 2 | 39877.70 | 39639.64 | 40115.86 | 3837.10 | 39849.50 | 4916.50 | 39661.68 | 40118.56 | Normal CI |
| Glaciale | 1 | 39625.40 | 39388.77 | 39861.97 | 3812.80 | 39597.30 | 4885.40 | 39410.66 | 39864.66 | Normal CI |
| Phaeophycae | 2 | 272646.90 | 251769.56 | 291612.39 | 314317.70 | 165427.50 | 301398.80 | 148286.16 | 181111.77 | Bootstrap CI |
| Phaeophycae | 1 | 280846.80 | 260565.11 | 300431.69 | 323770.70 | 170403.40 | 310464.80 | 152746.00 | 186559.58 | Bootstrap CI |
| Rhodophyta | 2 | 385345.50 | 352655.38 | 417965.42 | 514016.00 | 184221.40 | 446691.00 | 168621.39 | 209657.46 | Bootstrap CI |
| Rhodophyta | 1 | 398955.00 | 363452.97 | 432587.17 | 532169.50 | 190727.80 | 462466.70 | 174576.84 | 217062.29 | Bootstrap CI |
| Zostera Marina | 2 | 10183.80 | 9890.16 | 10477.40 | 4731.60 | 10097.40 | 6555.20 | 9636.49 | 10347.25 | Normal CI |
| Zostera Marina | 1 | 10135.80 | 9843.58 | 10428.05 | 4709.30 | 10049.70 | 6524.30 | 9591.15 | 10298.41 | Normal CI |
| Calcaereum | 2 | 47685.60 | 47379.10 | 47992.20 | 4940.00 | 47718.30 | 6996.90 | 47362.72 | 48310.01 | Normal CI |
| Calcaereum | 1 | 46119.70 | 45823.18 | 46416.15 | 4777.80 | 46151.20 | 6767.00 | 45807.38 | 46723.54 | Normal CI |
| Corallioides | 2 | 2987.10 | 2968.59 | 3005.53 | 297.70 | 2994.00 | 402.20 | 2968.18 | 3012.68 | Normal CI |
| Corallioides | 1 | 2987.10 | 2968.59 | 3005.53 | 297.70 | 2994.00 | 402.20 | 2968.18 | 3012.68 | Normal CI |
| Glaciale | 2 | 39879.50 | 39641.34 | 40117.58 | 3837.30 | 39851.60 | 4916.90 | 39663.69 | 40120.57 | Normal CI |
| Glaciale | 1 | 39627.10 | 39390.47 | 39863.69 | 3813.00 | 39599.40 | 4885.80 | 39412.67 | 39866.66 | Normal CI |
| Phaeophycae | 2 | 272651.90 | 252850.59 | 292499.72 | 314321.20 | 165427.50 | 301398.70 | 148286.41 | 181111.77 | Bootstrap CI |
| Phaeophycae | 1 | 280851.80 | 261170.18 | 300295.76 | 323774.20 | 170403.40 | 310464.70 | 152746.25 | 186559.58 | Bootstrap CI |
| Rhodophyta | 2 | 385346.60 | 352750.24 | 417014.98 | 514016.20 | 184221.40 | 446691.00 | 168621.39 | 209657.48 | Bootstrap CI |
| Rhodophyta | 1 | 398956.10 | 365408.67 | 432860.13 | 532169.70 | 190727.80 | 462466.70 | 174576.84 | 217062.31 | Bootstrap CI |
| Zostera Marina | 2 | 10191.50 | 9897.69 | 10485.28 | 4734.40 | 10097.40 | 6596.00 | 9660.60 | 10347.29 | Normal CI |
| Zostera Marina | 1 | 10143.50 | 9851.11 | 10435.93 | 4712.10 | 10049.70 | 6564.90 | 9614.99 | 10298.44 | Normal CI |
| Calcaereum | 2 | 47687.50 | 47380.93 | 47994.06 | 4940.20 | 47719.70 | 6995.60 | 47365.47 | 48312.22 | Normal CI |
| Calcaereum | 1 | 46121.50 | 45825.01 | 46418.00 | 4778.00 | 46152.70 | 6765.80 | 45810.14 | 46725.75 | Normal CI |
| Corallioides | 2 | 2988.90 | 2970.37 | 3007.34 | 297.90 | 2996.00 | 402.40 | 2969.71 | 3015.18 | Normal CI |
| Corallioides | 1 | 2988.90 | 2970.37 | 3007.34 | 297.90 | 2996.00 | 402.40 | 2969.71 | 3015.18 | Normal CI |
| Glaciale | 2 | 39881.30 | 39643.13 | 40119.39 | 3837.50 | 39853.50 | 4916.70 | 39665.74 | 40122.16 | Normal CI |
| Glaciale | 1 | 39628.90 | 39392.26 | 39865.51 | 3813.20 | 39601.30 | 4885.50 | 39414.72 | 39868.25 | Normal CI |
| Phaeophycae | 2 | 272659.10 | 251581.26 | 292807.49 | 314333.10 | 165427.50 | 301399.30 | 148340.35 | 181111.82 | Bootstrap CI |
| Phaeophycae | 1 | 280859.00 | 259530.57 | 300320.96 | 323786.20 | 170403.40 | 310465.30 | 152800.19 | 186559.63 | Bootstrap CI |
| Rhodophyta | 2 | 385348.10 | 353820.06 | 415130.13 | 514016.90 | 184221.60 | 446691.00 | 168621.42 | 209657.54 | Bootstrap CI |
| Rhodophyta | 1 | 398957.60 | 366427.76 | 430687.92 | 532170.40 | 190728.00 | 462466.70 | 174576.87 | 217062.37 | Bootstrap CI |
| Zostera Marina | 2 | 10197.50 | 9903.21 | 10491.70 | 4741.70 | 10097.40 | 6588.90 | 9699.86 | 10347.33 | Normal CI |
| Zostera Marina | 1 | 10149.50 | 9856.63 | 10442.35 | 4719.40 | 10049.70 | 6557.70 | 9656.06 | 10298.48 | Normal CI |
| Calcaereum | 2 | 47689.40 | 47382.84 | 47995.99 | 4940.40 | 47721.30 | 6996.30 | 47367.37 | 48314.30 | Normal CI |
| Calcaereum | 1 | 46123.40 | 45826.92 | 46419.94 | 4778.20 | 46154.30 | 6766.40 | 45812.03 | 46727.83 | Normal CI |
| Corallioides | 2 | 2990.80 | 2972.28 | 3009.27 | 298.10 | 2998.00 | 402.30 | 2971.77 | 3017.57 | Normal CI |
| Corallioides | 1 | 2990.80 | 2972.28 | 3009.27 | 298.10 | 2998.00 | 402.30 | 2971.77 | 3017.57 | Normal CI |
| Glaciale | 2 | 39883.20 | 39645.02 | 40121.30 | 3837.60 | 39855.70 | 4917.10 | 39668.34 | 40123.75 | Normal CI |
| Glaciale | 1 | 39630.80 | 39394.14 | 39867.42 | 3813.40 | 39603.50 | 4886.00 | 39417.32 | 39869.84 | Normal CI |
| Phaeophycae | 2 | 272669.10 | 251755.75 | 291442.14 | 314341.70 | 165427.50 | 301399.40 | 148340.36 | 181111.98 | Bootstrap CI |
| Phaeophycae | 1 | 280869.00 | 259646.52 | 300622.87 | 323794.80 | 170403.40 | 310465.40 | 152800.20 | 186559.79 | Bootstrap CI |
| Rhodophyta | 2 | 385349.90 | 352075.55 | 417067.40 | 514021.20 | 184221.70 | 446691.10 | 168621.42 | 209657.58 | Bootstrap CI |
| Rhodophyta | 1 | 398959.30 | 364010.78 | 432962.46 | 532174.70 | 190728.10 | 462466.80 | 174576.88 | 217062.41 | Bootstrap CI |
| Zostera Marina | 2 | 10202.60 | 9908.35 | 10496.89 | 4742.10 | 10097.50 | 6585.30 | 9719.36 | 10348.38 | Normal CI |
| Zostera Marina | 1 | 10154.70 | 9861.76 | 10447.54 | 4719.90 | 10049.80 | 6554.10 | 9673.49 | 10299.53 | Normal CI |
| Calcaereum | 2 | 47691.40 | 47384.81 | 47997.98 | 4940.60 | 47723.30 | 6996.70 | 47369.15 | 48316.56 | Normal CI |
| Calcaereum | 1 | 46125.40 | 45828.89 | 46421.93 | 4778.30 | 46156.20 | 6766.80 | 45813.81 | 46730.09 | Normal CI |
| Corallioides | 2 | 2992.80 | 2974.26 | 3011.28 | 298.30 | 3000.20 | 401.90 | 2974.17 | 3019.27 | Normal CI |
| Corallioides | 1 | 2992.80 | 2974.26 | 3011.28 | 298.30 | 3000.20 | 401.90 | 2974.17 | 3019.27 | Normal CI |
| Glaciale | 2 | 39885.20 | 39647.02 | 40123.33 | 3837.80 | 39858.30 | 4917.70 | 39669.83 | 40124.99 | Normal CI |
| Glaciale | 1 | 39632.80 | 39396.15 | 39869.45 | 3813.60 | 39606.10 | 4886.60 | 39418.81 | 39871.09 | Normal CI |
| Phaeophycae | 2 | 272677.00 | 253175.97 | 290339.80 | 314351.70 | 165427.50 | 301399.20 | 148340.36 | 181111.98 | Bootstrap CI |
| Phaeophycae | 1 | 280876.90 | 260178.27 | 301353.79 | 323804.70 | 170403.50 | 310465.20 | 152800.20 | 186559.79 | Bootstrap CI |
| Rhodophyta | 2 | 385353.80 | 353296.56 | 416578.31 | 514025.40 | 184221.70 | 446691.20 | 168621.42 | 209657.58 | Bootstrap CI |
| Rhodophyta | 1 | 398963.20 | 364098.03 | 433600.97 | 532178.90 | 190728.10 | 462466.90 | 174576.88 | 217062.41 | Bootstrap CI |
| Zostera Marina | 2 | 10210.10 | 9915.42 | 10504.76 | 4748.60 | 10097.50 | 6600.70 | 9722.58 | 10348.42 | Normal CI |
| Zostera Marina | 1 | 10162.10 | 9868.83 | 10455.42 | 4726.30 | 10049.90 | 6569.50 | 9677.03 | 10299.56 | Normal CI |
| Calcaereum | 2 | 47693.50 | 47386.92 | 48000.11 | 4940.80 | 47725.60 | 6998.80 | 47371.73 | 48319.48 | Normal CI |
| Calcaereum | 1 | 46127.50 | 45831.00 | 46424.06 | 4778.50 | 46158.70 | 6768.90 | 45816.39 | 46733.01 | Normal CI |
| Corallioides | 2 | 2994.90 | 2976.35 | 3013.40 | 298.50 | 3002.30 | 402.10 | 2975.34 | 3021.23 | Normal CI |
| Corallioides | 1 | 2994.90 | 2976.35 | 3013.40 | 298.50 | 3002.30 | 402.10 | 2975.34 | 3021.23 | Normal CI |
| Glaciale | 2 | 39887.30 | 39649.09 | 40125.43 | 3838.00 | 39859.80 | 4916.70 | 39671.59 | 40127.04 | Normal CI |
| Glaciale | 1 | 39634.90 | 39398.22 | 39871.54 | 3813.80 | 39607.50 | 4885.60 | 39420.57 | 39873.13 | Normal CI |
| Phaeophycae | 2 | 272679.40 | 251881.50 | 293143.51 | 314351.60 | 165427.50 | 301399.20 | 148340.36 | 181115.10 | Bootstrap CI |
| Phaeophycae | 1 | 280879.30 | 260866.52 | 299281.96 | 323804.60 | 170403.50 | 310465.20 | 152800.20 | 186562.91 | Bootstrap CI |
| Rhodophyta | 2 | 385354.90 | 355014.61 | 418585.78 | 514027.20 | 184221.70 | 446692.00 | 168621.42 | 209657.61 | Bootstrap CI |
| Rhodophyta | 1 | 398964.30 | 364385.42 | 431311.67 | 532180.70 | 190728.10 | 462467.70 | 174576.88 | 217062.44 | Bootstrap CI |
| Zostera Marina | 2 | 10214.90 | 9919.84 | 10509.94 | 4754.70 | 10097.90 | 6600.10 | 9722.59 | 10348.60 | Normal CI |
| Zostera Marina | 1 | 10166.90 | 9873.25 | 10460.59 | 4732.40 | 10050.30 | 6568.90 | 9677.04 | 10299.76 | Normal CI |
| Calcaereum | 2 | 47695.70 | 47389.13 | 48002.35 | 4941.00 | 47728.50 | 6998.30 | 47373.54 | 48321.10 | Normal CI |
| Calcaereum | 1 | 46129.80 | 45833.21 | 46426.30 | 4778.70 | 46161.50 | 6768.50 | 45818.21 | 46734.63 | Normal CI |
| Corallioides | 2 | 2997.10 | 2978.60 | 3015.68 | 298.80 | 3004.20 | 402.20 | 2977.52 | 3023.77 | Normal CI |
| Corallioides | 1 | 2997.10 | 2978.60 | 3015.68 | 298.80 | 3004.20 | 402.20 | 2977.52 | 3023.77 | Normal CI |
| Glaciale | 2 | 39889.50 | 39651.33 | 40127.70 | 3838.30 | 39861.60 | 4917.50 | 39675.34 | 40128.98 | Normal CI |
| Glaciale | 1 | 39637.10 | 39400.45 | 39873.81 | 3814.00 | 39609.30 | 4886.30 | 39424.32 | 39875.06 | Normal CI |
| Phaeophycae | 2 | 272684.20 | 250944.48 | 291410.17 | 314355.80 | 165427.50 | 301399.20 | 148340.37 | 181115.31 | Bootstrap CI |
| Phaeophycae | 1 | 280884.10 | 258668.08 | 300583.02 | 323808.80 | 170403.50 | 310465.20 | 152800.21 | 186563.12 | Bootstrap CI |
| Rhodophyta | 2 | 385355.50 | 352816.18 | 415372.30 | 514027.50 | 184221.70 | 446692.00 | 168621.42 | 209657.61 | Bootstrap CI |
| Rhodophyta | 1 | 398965.00 | 364337.82 | 429727.21 | 532181.00 | 190728.10 | 462467.70 | 174576.88 | 217062.44 | Bootstrap CI |
| Zostera Marina | 2 | 10220.90 | 9925.76 | 10515.96 | 4755.50 | 10098.00 | 6566.30 | 9722.61 | 10349.29 | Normal CI |
| Zostera Marina | 1 | 10172.90 | 9879.18 | 10466.62 | 4733.20 | 10050.30 | 6535.30 | 9677.06 | 10300.44 | Normal CI |
| Calcaereum | 2 | 47698.10 | 47391.44 | 48004.69 | 4941.20 | 47731.80 | 6999.00 | 47375.26 | 48324.39 | Normal CI |
| Calcaereum | 1 | 46132.10 | 45835.52 | 46428.64 | 4779.00 | 46164.80 | 6769.10 | 45819.93 | 46737.92 | Normal CI |
| Corallioides | 2 | 2999.50 | 2980.94 | 3018.05 | 299.00 | 3006.70 | 402.20 | 2979.30 | 3026.24 | Normal CI |
| Corallioides | 1 | 2999.50 | 2980.94 | 3018.05 | 299.00 | 3006.70 | 402.20 | 2979.30 | 3026.24 | Normal CI |
| Glaciale | 2 | 39891.80 | 39653.63 | 40130.03 | 3838.50 | 39863.10 | 4917.60 | 39678.25 | 40130.81 | Normal CI |
| Glaciale | 1 | 39639.40 | 39402.76 | 39876.14 | 3814.20 | 39610.90 | 4886.50 | 39427.23 | 39876.90 | Normal CI |
| Phaeophycae | 2 | 272691.30 | 252482.75 | 292095.14 | 314359.60 | 165427.50 | 301399.20 | 148345.57 | 181115.32 | Bootstrap CI |
| Phaeophycae | 1 | 280891.20 | 259084.57 | 300981.07 | 323812.60 | 170403.50 | 310465.20 | 152805.41 | 186563.12 | Bootstrap CI |
| Rhodophyta | 2 | 385356.70 | 353413.27 | 418639.00 | 514028.70 | 184221.70 | 446692.10 | 168621.43 | 209658.25 | Bootstrap CI |
| Rhodophyta | 1 | 398966.10 | 364346.02 | 431744.86 | 532182.20 | 190728.10 | 462467.80 | 174576.88 | 217063.08 | Bootstrap CI |
| Zostera Marina | 2 | 10223.90 | 9928.79 | 10519.09 | 4756.30 | 10099.40 | 6570.60 | 9722.61 | 10349.40 | Normal CI |
| Zostera Marina | 1 | 10176.00 | 9882.20 | 10469.75 | 4734.10 | 10051.70 | 6539.70 | 9677.90 | 10300.54 | Normal CI |
| Total | 2 | 758614.30 | 722564.40 | 795742.37 | 594629.60 | 573049.70 | 640534.60 | 533206.60 | 610521.20 | bootstrap CI |
| Total | 1 | 778557.30 | 740114.79 | 815633.90 | 614798.20 | 587123.00 | 663186.40 | 545460.40 | 625558.40 | bootstrap CI |
| Total | 2 | 758626.80 | 720298.99 | 792748.01 | 594641.90 | 573052.90 | 640529.80 | 533211.20 | 610523.80 | bootstrap CI |
| Total | 1 | 778569.80 | 739898.39 | 818046.72 | 614810.40 | 587126.20 | 663181.60 | 545463.60 | 625562.10 | bootstrap CI |
| Total | 2 | 758637.50 | 718217.24 | 794053.15 | 594645.00 | 573057.20 | 640645.30 | 533214.90 | 610769.00 | bootstrap CI |
| Total | 1 | 778580.50 | 736521.44 | 818777.57 | 614813.60 | 587130.40 | 663297.10 | 545467.50 | 625565.50 | bootstrap CI |
| Total | 2 | 758647.90 | 720455.44 | 796470.50 | 594646.20 | 573061.70 | 640653.10 | 533218.60 | 610771.90 | bootstrap CI |
| Total | 1 | 778590.90 | 744905.60 | 816060.60 | 614814.80 | 587134.90 | 663304.90 | 545472.10 | 625570.40 | bootstrap CI |
| Total | 2 | 758666.00 | 717826.32 | 796498.52 | 594656.70 | 573085.80 | 640646.30 | 533221.90 | 610774.70 | bootstrap CI |
| Total | 1 | 778609.00 | 740664.62 | 818653.89 | 614825.20 | 587159.10 | 663298.10 | 545491.90 | 625573.70 | bootstrap CI |
| Total | 2 | 758682.60 | 719709.46 | 795149.65 | 594662.90 | 573090.60 | 640686.20 | 533226.10 | 610778.90 | bootstrap CI |
| Total | 1 | 778625.60 | 739144.51 | 817999.39 | 614831.50 | 587163.90 | 663338.00 | 545498.00 | 625580.70 | bootstrap CI |
| Total | 2 | 758703.30 | 721391.60 | 794266.81 | 594672.40 | 573096.20 | 640684.10 | 533238.40 | 610782.80 | bootstrap CI |
| Total | 1 | 778646.30 | 740615.84 | 816406.52 | 614840.90 | 587169.40 | 663335.90 | 545502.60 | 625585.90 | bootstrap CI |
| Total | 2 | 758723.30 | 720156.96 | 795857.17 | 594676.80 | 573100.10 | 640682.90 | 533243.50 | 610786.30 | bootstrap CI |
| Total | 1 | 778666.20 | 741664.20 | 816401.32 | 614845.30 | 587173.30 | 663334.70 | 545508.80 | 625689.60 | bootstrap CI |
| Total | 2 | 758742.20 | 720920.60 | 795009.88 | 594676.10 | 573131.30 | 640719.40 | 533251.70 | 610796.50 | bootstrap CI |
| Total | 1 | 778685.20 | 741017.49 | 816170.91 | 614844.60 | 587204.60 | 663371.30 | 545513.10 | 625695.20 | bootstrap CI |
| Total | 2 | 758762.30 | 724353.26 | 796817.79 | 594677.90 | 573137.40 | 640718.50 | 533257.80 | 610801.10 | bootstrap CI |
| Total | 1 | 778705.30 | 738927.11 | 814183.65 | 614846.40 | 587210.70 | 663370.30 | 545517.40 | 625700.40 | bootstrap CI |
| Total | 2 | 758784.90 | 720292.42 | 798803.94 | 594687.10 | 573143.30 | 640715.40 | 533262.60 | 610805.80 | bootstrap CI |
| Total | 1 | 778727.90 | 739600.00 | 815544.32 | 614855.50 | 587216.60 | 663367.20 | 545541.50 | 625705.50 | bootstrap CI |
| Total | 2 | 758810.20 | 722729.86 | 793640.17 | 594694.00 | 574592.40 | 640377.20 | 533267.60 | 610813.80 | bootstrap CI |
| Total | 1 | 778753.20 | 739635.97 | 822067.24 | 614862.50 | 588665.60 | 663161.80 | 545547.40 | 625711.50 | bootstrap CI |
| Total | 2 | 758824.90 | 721793.54 | 794066.05 | 594696.90 | 574621.20 | 640376.50 | 533272.40 | 610820.90 | bootstrap CI |
| Total | 1 | 778767.90 | 741563.78 | 815064.85 | 614865.30 | 588694.40 | 663161.20 | 545553.90 | 625717.80 | bootstrap CI |
| Total | 2 | 758843.00 | 722051.04 | 796558.80 | 594699.20 | 574784.50 | 640375.50 | 533282.50 | 611031.00 | bootstrap CI |
| Total | 1 | 778786.00 | 740039.40 | 819916.33 | 614867.70 | 588996.20 | 663160.10 | 545559.10 | 625726.30 | bootstrap CI |
| Total | 2 | 758861.30 | 719510.29 | 796259.36 | 594701.60 | 574790.90 | 640375.30 | 533290.50 | 611044.50 | bootstrap CI |
| Total | 1 | 778804.30 | 740891.45 | 817866.23 | 614870.10 | 589002.60 | 663160.00 | 545568.70 | 625732.10 | bootstrap CI |

**Table S11.** Mann-Whitney U test comparing sequestration rates across Scenarios in 2026 and 2040. p-values adjusted using the Benjamini–Hochberg correction for multiple comparisons

| **Species** | **CalendarYear** | **p_value** | **U_statistic** | **p_adj** | **Significant** |
| --- | --- | --- | --- | --- | --- |
| Calcaereum | 2026 | 3.54E-12 | 5.90E+05 | 2.15E-11 | Yes |
| Calcaereum | 2040 | 3.58E-12 | 5.90E+05 | 2.15E-11 | Yes |
| Corallioides | 2026 | 1.00E+00 | 5.00E+05 | 1.00E+00 | No |
| Corallioides | 2040 | 1.00E+00 | 5.00E+05 | 1.00E+00 | No |
| Glaciale | 2026 | 1.34E-01 | 5.19E+05 | 4.02E-01 | No |
| Glaciale | 2040 | 1.34E-01 | 5.19E+05 | 4.02E-01 | No |
| Phaeophycae | 2026 | 5.87E-01 | 4.93E+05 | 8.97E-01 | No |
| Phaeophycae | 2040 | 5.87E-01 | 4.93E+05 | 8.97E-01 | No |
| Rhodophyta | 2026 | 5.98E-01 | 4.93E+05 | 8.97E-01 | No |
| Rhodophyta | 2040 | 5.98E-01 | 4.93E+05 | 8.97E-01 | No |
| Zostera Marina | 2026 | 7.97E-01 | 5.03E+05 | 9.57E-01 | No |
| Zostera Marina | 2040 | 7.98E-01 | 5.03E+05 | 9.57E-01 | No |

**Table S12.** Sequestration in 2040 (t C yr^-1^) from restoration projects developed from 2026-2040

| **Species** | **Scenario** | **Median_Delta** | **Lower_CI** | **Upper_CI** | **Mean_Delta** | **SD_Delta** |
| --- | --- | --- | --- | --- | --- | --- |
| Calcaereum | 2 | 23.68 | 23.24 | 24.15 | 24.02 | 5.38 |
| Calcaereum | 1 | 23.68 | 23.24 | 24.15 | 24.02 | 5.38 |
| Corallioides | 2 | 23.35 | 22.86 | 23.76 | 23.91 | 5.22 |
| Corallioides | 1 | 23.35 | 22.86 | 23.76 | 23.91 | 5.22 |
| Glaciale | 2 | 23.06 | 22.78 | 23.55 | 23.79 | 5.45 |
| Glaciale | 1 | 23.06 | 22.78 | 23.55 | 23.79 | 5.45 |
| Phaeophycae | 2 | 7.58 | 6.17 | 9.21 | 86.48 | 311.30 |
| Phaeophycae | 1 | 7.58 | 6.17 | 9.21 | 86.48 | 311.30 |
| Rhodophyta | 2 | 1.45 | 1.19 | 1.76 | 24.47 | 110.22 |
| Rhodophyta | 1 | 1.45 | 1.19 | 1.76 | 24.47 | 110.22 |
| Zostera Marina | 2 | 9.57 | 8.21 | 10.77 | 64.35 | 257.08 |
| Zostera Marina | 1 | 9.57 | 8.21 | 10.77 | 64.35 | 257.08 |
| Total | 2 | 115.00 | 110.87 | 122.32 | 247.03 | 415.59 |
| Total | 1 | 115.00 | 110.87 | 122.32 | 247.03 | 415.59 |

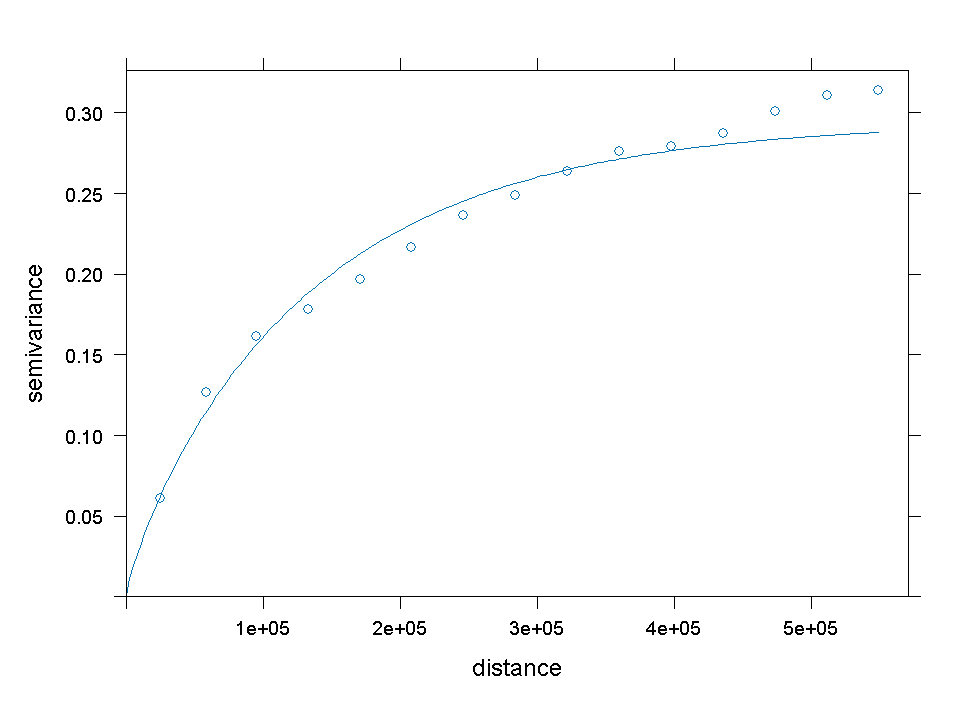
**Figure S1.** Empirical and fitted variogram for log-transformed seabed flow velocity using depth as a covariate

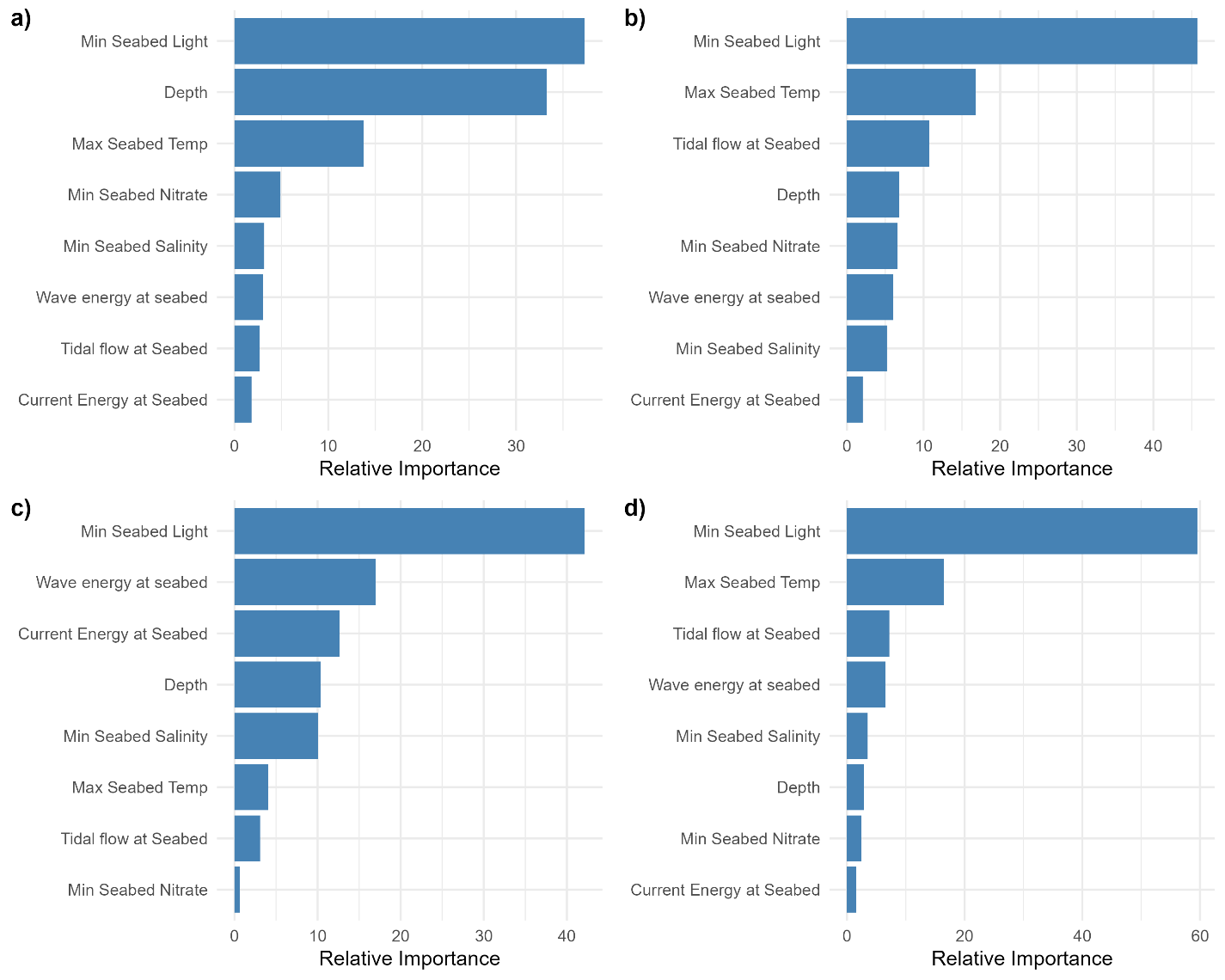
**Figure S2.** Relative feature importance for the boosted regression tree (BRT) species distribution models for (a) *Zostera marina*, (b) *Phymatolithon calcareum*, (c*) Lithothamnion corallioides*, and (d) *Lithothamnion glaciale*. Relative importance is the average of the number of splits of the variable calculated based on the number of times a variable is selected for splitting, weighted by the squared improvement to the model due to the split

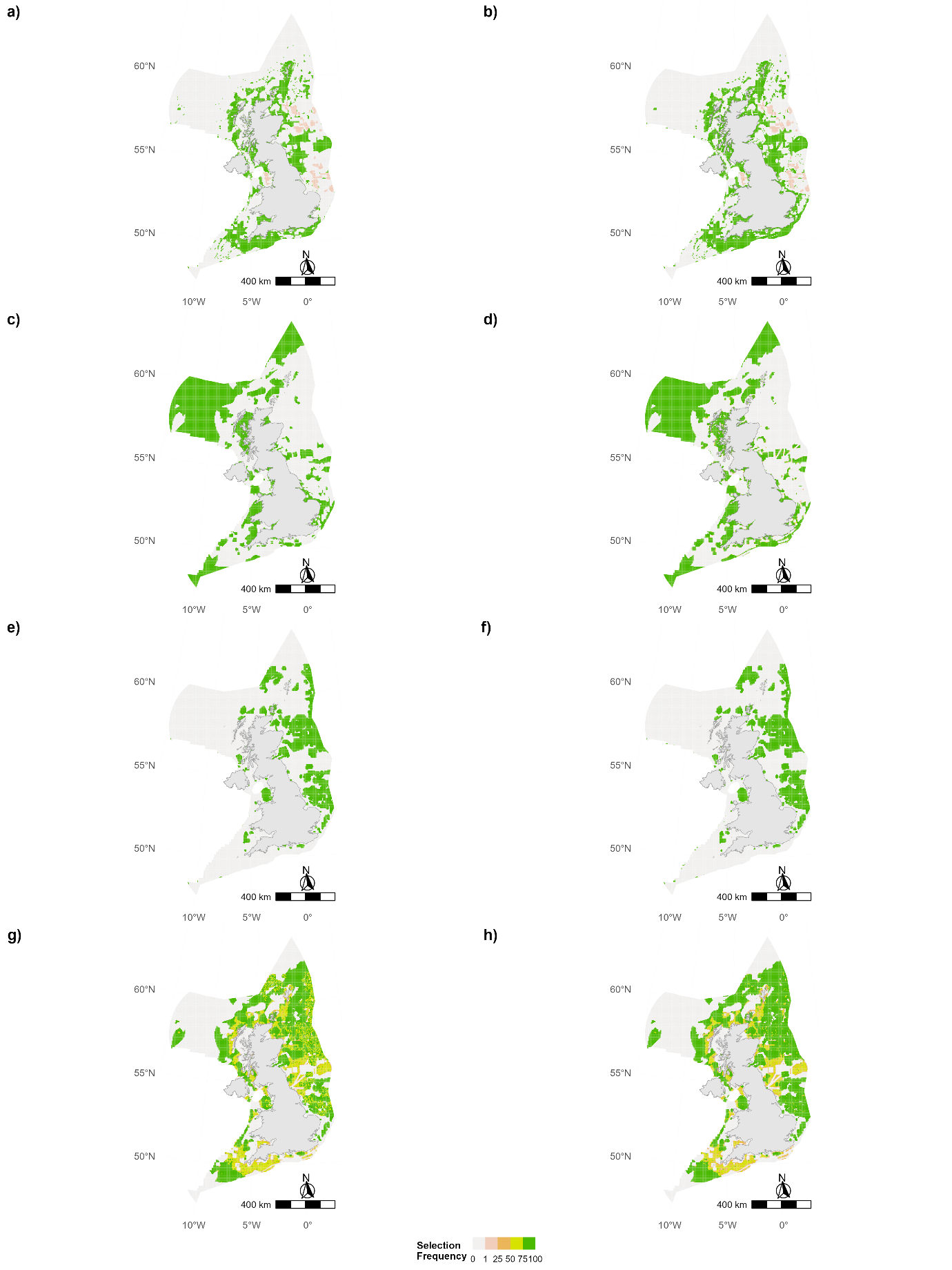
**Figure S3.** Comparative analysis of spatial unit selection frequency for offshore activities across two scenarios for 1000 runs a) Scenario 1 Open b) Scenario 2 Open c) Scenario 1 Conservation d) Scenario 2 Conservation e) Scenario 1 Oil,f) Scenario 2 Oil g) Scenario 1 Energy h) Scenario 2 Energy. Higher frequencies suggest more consistent selection of these units under the respective scenario and use type

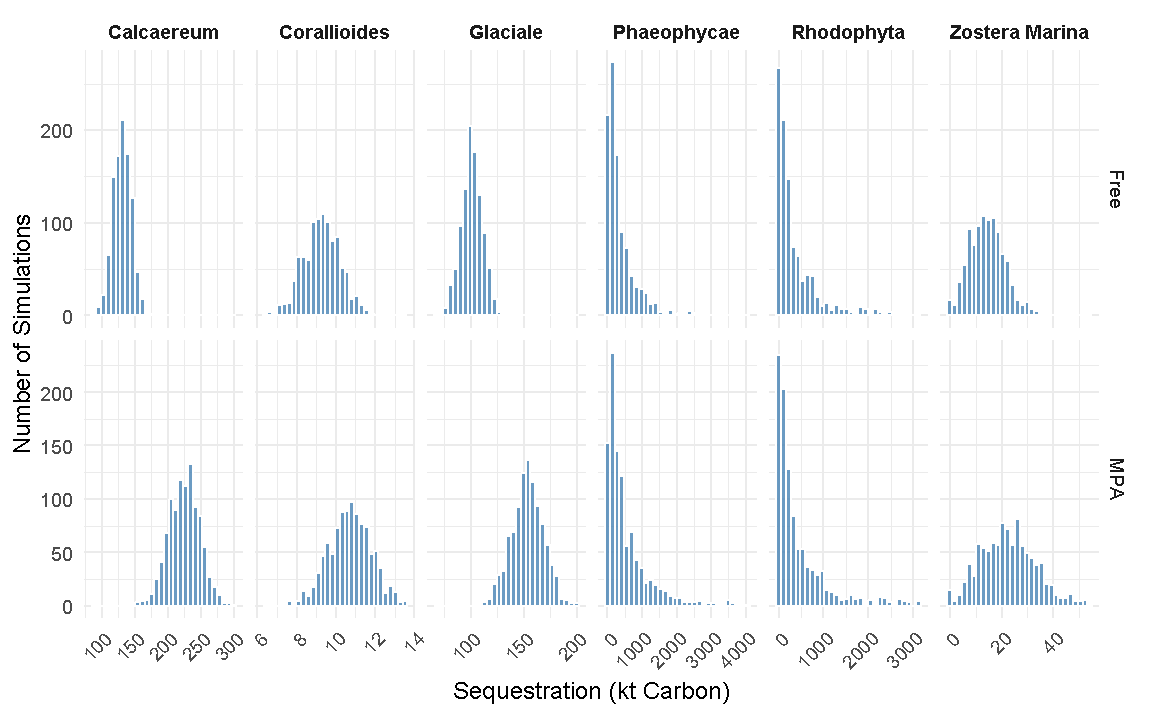
**Figure S4.** Blue carbon sequestration potentials in 2040 (kt C yr⁻¹) from 1,000 Monte Carlo simulations under Scenario 1 (MPA) and Scenario 2 (Free). (a) *Zostera marina*, (b) Rhodophyta spp., (c) Phaeophyceae spp., (d) *Phymatolithon calcareum*, (e) *Lithothamnion corallioides*, and (f) *Lithothamnion glaciale*.
